# Climatic niche conservatism and ecological diversification in the Holarctic cold-dwelling butterfly genus *Erebia*

**DOI:** 10.1101/2022.04.12.488065

**Authors:** Irena Klečková, Jan Klečka, Zdeněk Faltýnek Fric, Martin Česánek, Ludovic Dutoit, Loïc Pellissier, Pável Matos-Maraví

## Abstract

The diversification of alpine species has been modulated by their climatic niches interacting with changing climatic conditions. The relative roles of climatic niche conservatism promoting geographical speciation and of climatic niche diversification are poorly understood in diverse temperate groups. Here, we investigate the climatic niche evolution in a species rich butterfly genus, *Erebia*. This Holarctic cold-dwelling genus reaches the highest diversity in European mountains. We generated a nearly complete molecular phylogeny and modelled the climatic niche evolution using geo-referenced occurrence records. We reconstructed the evolution of the climatic niche and tested how the species’ climatic niche width changes across the occupied climate gradient and compared two main *Erebia* clades, the European and the Asian clade. We further explored climatic niche overlaps among species. Our analyses revealed that the evolution of *Erebia* has been shaped by climatic niche conservatism, supported by a strong phylogenetic signal and niche overlap in sister species, likely promoting allopatric speciation. The European and the Asian clades evolved their climatic niches toward different local optima. In addition, species in the European clade have narrower niches compared to the Asian clade. Contrasts among the clades may be related to regional climate differences, with lower climate seasonality in Europe compared to Central Asia favouring the evolution of narrower niches. Further, adaptive divergence could appear in other traits, such as habitat use, which can be reflected by narrower climatic niches detected in the European clade. In conclusion, our study extends knowledge about the complexity of evolutionary drivers in temperate insects.

## INTRODUCTION

Mountains are major global biodiversity reservoirs whose communities have been shaped by evolutionary and ecological processes linked with topographical complexity and elevational gradients (Wiens and Graham 2005, Schmitt et al. 2016, Condamine et al. 2018). Temperate mountains generally host lower species diversity compared to tropical mountains. The climatic niche, which is primarily defined by temperature and precipitation regimes, modulates where species occur and how they respond to climatic changes (Deutsch et al. 2008, Bonetti and Wiens 2014). In the tropics, narrower climatic niches (Janzen 1967) and rapid climatic niche evolution (lability) might have further accelerated diversification rates and driven a greater accumulation of taxa (Kozak and Wiens 2010). In contrast, a pattern of climatic niche conservatism associated with broad temperature seasonality may have dominated in the diversification of temperate lineages (Kozak and Wiens 2007). The climatic niche conservatism shapes regional species diversity by dispersal along elevational gradients and across valleys during phases of climatic driven range expansions (Cadena et al. 2012, Qian et al. 2021), potentially driving allopatric speciation following range contractions (Xing and Ree 2017, Ding et al. 2020). While niche conservatism associated with broad climatic niches may have determined the extant species richness in several temperate groups of vertebrates (Smith et al. 2005, Pyron and Burbrink 2009), the role of macro-evolutionary mechanisms shaping lineage diversity in temperate mountains should be assessed in invertebrates.

The extant species diversity of temperate mountains has been primarily shaped by dispersal mediated by climatic niche conservatism (Condamine et al. 2018, Liu et al. 2021), but also by specialisation to extreme climatic conditions (Su et al. 2020), and by diversity-dependent radiation across empty niches on isolated mountain ranges (Peña et al. 2015). Diversity-dependent radiation might have resulted from various inter-specific interactions (e.g. interactions with host plants or competitive exclusion among closely related butterfly species) in diverse groups promoting diversification until a macroevolutionary saturation point is reached and species diversity is maintained through time (Calcagno et al. 2017). Sympatric speciation, which leads to differentiation of species with overlapping ranges, can be also driven by climatic niche conservatism or niche divergence (Pitteloud et al. 2017), which may be promoted along climatic gradients across mountain slopes. At climatic margins of occurrence, a decrease of fitness may enhance species diversification by niche specialisation as an adaptive response to the extreme climates (O’Brien et al. 2022). However, the role of niche specialisation across climatic gradients as a possible driver of species diversification is still understudied.

The butterfly genus *Erebia* is a cold-dwelling temperate group with high species diversity. The genus contains 90-100 species (Warren 1936, Tennent 2008), and most of its diversity is concentrated in mountain ranges across the Holarctic region, with several species occurring in the Arctic and a few species found in lowlands (Warren 1936). *Erebia* larvae are mainly polyphagous, feeding on widespread grasses, sedges, and rushes (*Poaceae, Cyperaceae*, and *Juncaceae*) (Sonderegger, 2005). In captivity, different European species consume *Poa annua, Poa pratensis* or *Festuca rubra* (Sonderegger, 2005), but local populations may differ in their host plant use (Gunson 2019). Thus, the geographical distribution has likely not been primarily constrained by the host plants unlike in more specialised herbivorous insects (Su et al. 2020). An adaptive radiation scenario has been proposed for *Erebia* based on an apparent slowdown of diversification rates toward the present (Martin et al. 2000, Peña et al. 2015). Alternatively, a diversification scenario under climatic niche conservatism was invoked to explain dispersal during climatic fluctuations followed by vicariant differentiation (Schmitt et al. 2016). The adaptive scenario is supported by differences in optimal altitudes of occurrence in closely related species across mountain slopes in European Alps (Lorkovic 1958, Sonderegger 2005, Varga 2014) and by contrasting thermoregulation strategies by microhabitat use observed in sympatric butterflies co-occurring on a slope in European Alps (Kleckova et al. 2014). As mountain specialists, *Erebia* species living in proximity may inhabit areas with very different climates. For example, sites located above the treeline are significantly colder than sites located in the valleys a few kilometres away. Temperature and precipitation regimes may also substantially differ between south- and north-facing slopes in the same mountain range (Scherrer & Kőrner, 2010). The divergence in climatic traits can also reflect divergence in other traits such as habitat use. For example, coexistence of closely related species at the regional scale may be accompanied by competitive exclusion along the altitudinal gradient at the local scale (Lorkovic 1958).

Here, we ask whether the evolutionary history of a species-rich group of temperate cold-dwelling butterflies is determined by niche conservatisms or rather by adaption to distinct climatic niches among lineages. Studies of *Erebia* secondary contact zones suggest preceding allopatric differentiation connected with coevolution with endosymbionts (Lucek et al. 2021). To shed more light on processes driving the diversification of *Erebia* at the macroevolutionary and macroecological scales, we ask three sets of questions in this study:

1. Did species’ climatic niches evolve gradually following the niche conservatism hypothesis or rapidly during the early evolution of the genus in congruence with the adaptive radiation scenario? Did the climatic niches of different clades evolve toward different local optima?
2. Do sister species display more similar climatic niches compared to non-sister species pairs as expected under the niche conservatism scenario? Alternatively, do sympatric sister species have distinct climatic niches to reduce inter-specific competition?
3. Is the high species diversity found in European mountains explained by the accumulation of taxa with narrow climatic niches? Additionally, do species occurring in extreme climates on the periphery of their distributional ranges have broad climatic niches or are they climatic niche specialists?

## METHODS

### Phylogeny and divergence times

To obtain a nearly complete phylogeny of the genus *Erebia*, we extended the dataset from Peña et al. (2015) by sequencing 1 mitochondrial (COI) and 3 nuclear gene fragments (GAPDH, RpS5, wingless). Our dataset includes 8 outgroups and 83 representatives (one per species) of the genus *Erebia* (Supplementary materials, Table S1), covering 80-90% of the global species diversity of *Erebia*. Divergence times were estimated in the software BEAST v 2.6.0 (Bouckaert et al. 2019).

Alignment partitions and substitution models were specified by PartitionFinder 2 (Lanfear et al. 2012). However, due to low effective sample sizes (ESS) in preliminary analyses, we reduced the complexity of the substitution models for the mitochondrial partitions (Supplementary materials, Table S2). We used a log-normal relaxed molecular clock and the birth-death incomplete sampling process. Other priors were left as default. As fossil records have not been identified for the genus *Erebia*, we used secondary calibrations from a comprehensive, fossil based Nymphalidae chronogram (Chazot et al. 2021):

1. The root age of the tribe Satyrini Boisduval 1833 was defined by a uniform distribution between 37.45 to 45.55 Ma.
2. The crown node of a strongly supported clade within the tribe Satyrini consisting of the subtribes Satyrina and Melanargiina was specified by a uniform distribution between 23.12 to 31.39 Ma.
3. The split between the genera *Maniola* and *Erebia* had a uniform distribution between 26.98 to 35.64 Ma.

To evaluate our conservative secondary calibration strategy that favours non-informative prior distribution in the three constrained nodes, we used more informative normal prior distributions (Supplementary Material S1). For the subsequent analyses, we used the phylogenetic tree inferred by using uniform distribution of calibration points.

The length of the Markov chain Monte Carlo (MCMC) chain was set to 100 million steps, storing the estimated parameters every 10,000 steps. We reconstructed the consensus tree from 1000 randomly sampled trees from the posterior distribution using the program TreeAnnotator implemented in BEAST v1.10.4. software (Suchard et al. 2018) and calculated mean node heights to obtain divergence times. For downstream biogeographical and climatic niche evolution analyses, we pruned the outgroups in the estimated phylogeny by using the drop.tip() command in the ape package v. 5.6-2 (Paradis and Schliep 2019) using the software R v 4.1.2 (R Core Team 2021).

### Biogeography

We specified 3 biogeographical regions: Western Palaearctic (A), Eastern Palaearctic (B), and North America (C) (Supplementary materials, Table S3). We estimated ancestral ranges along the phylogeny of *Erebia* using the R package BioGeoBEARS v1.1.2 (Matzke 2013). We compared six competing biogeographical models using the Akaike information criterion (AICc) (Akaike 1973). The analysed models were the dispersal–extinction–cladogenesis model (DEC) (Ree and Smith 2008), a model resembling the dispersal-vicariance model of Ronquist (1997) (“DIVALIKE”) and a model resembling the BayArea model (Landis et al. 2013) (“BAYAREALIKE”). We also tested a modified version of each of these three models to include the founder-event speciation parameter J (Matzke 2014). Although the usage of the J parameter has been criticized (Ree and Sanmartín 2018), it has been recently shown that a comparison of models including such a parameter is statistically valid (Matzke 2021).

### Climatic data

We collected 25,694 occurrence points from the literature and biodiversity databases (Supplementary material, Table S4). For each species, we extracted bioclimatic variables from WorldClim v. 2.1. at the 30” resolution (about 1 km^2^, ESRI grid) in the R package Raster v. 3.5 (Hijmans and van Etten 2012). The environmental conditions where *Erebia* butterflies occur were described by four bioclimatic variables: annual mean temperature (bio1), temperature seasonality (bio4), annual precipitation (bio12), and precipitation seasonality (bio15). We selected these variables because they are known to affect species abundance and survival in *Erebia*. For example, the annual mean temperature affects the length of larval development, flight season and overwintering conditions, while the annual precipitation (reflecting humidity of the environment) drove *Erebia* distribution during the Pleistocene (Schmitt et al., 2006). Further, temperature seasonality and precipitation seasonality describe fluctuations of the variables during the year and can be used as a proxy for climatic extremes affecting *Erebia* population abundances (Konvicka et al. 2021, Vrba et al. 2017). Larger values of seasonality indices represent larger variability of temperature or precipitation during the year (O’Donnel and Ignizio 2012). Further, we extracted the WorldClim variable Elevation for every occurrence point and used the mean elevation of species occurrence records as an additional proxy for the climatic niche. We then used Principal Component Analysis (PCA) and duality diagrams (dudi) implemented in R package ade4 v. 1.7.18 (Dray and Dufour 2007) to visualise the distribution-based climatic data for individuals of all species and to compare the overall climatic niche of the Asian and the European clades.

### Evolution of climatic traits

First, we computed the average value of each bioclimatic variable and elevation for each species and used them as climatic traits (e.g., Arnal et al., 2019). Then, we compared the fit of three models to describe the evolution of the climatic traits and elevation: the Brownian Motion (BM), the Ornstein-Uhlenbeck (OU) (Butler and King 2004) and the Early Burst (EB) models (Simpson 1953). We used the R package geiger v. 2.0.10 (Pennell et al. 2014) and its fitContinuous() function to compare model fit according to AICc. Under Brownian motion, the values of the evolving trait are expected to change randomly, without limits, along branches in a phylogenetic tree. The Brownian motion model includes the Brownian rate parameter (σ^2^), which describes how fast the traits randomly change through time. Under the OU model, a trait is expected to evolve toward a local optimum. The OU model includes the parameter θ describing the optimal value of the trait (long term mean value of the trait), σ^2^ as in the BM, and the attraction strength α (Cooper et al. 2016). If α = 0, the OU model is identical with BM, and higher values of α suggest strong deviation from a simple random walk and a faster evolution toward the optimum trait value. Under the Early Burst model, consistent with an adaptive radiation scenario, it is expected that the rate of trait evolution, described by the parameter *a*, decreases trough time, having the highest value close to the root of the phylogeny.

To quantitatively measure the degree to which evolutionary history predicts ecological similarity among species, we tested whether there is a phylogenetic signal in the mean values of the bioclimatic variables and the elevation. The recovery of a phylogenetic signal is one of several indicators of trait conservatism as opposed to trait lability, i.e., it indicates that closely related species share more similar trait states than random taxa from the phylogeny (Blomberg et al. 2003). Under the adaptive radiation scenario, implying a higher importance of species interactions (e.g., competition for resources), it is expected that the climatic niches among sister species evolve rapidly, potentially obscuring any phylogenetic signal, whereas under the niche conservatism scenario, the presence of a phylogenetic signal would indicate that sister species conserve their climatic niche optima and differentiation might have been driven by other mechanisms, e.g., allopatric speciation. We used the phylosig() function of the R package Phytools v. 0.7.90 (Revell 2012) to calculate two indices of phylogenetic signal, Pagel’s λ (Pagel 1999) and Blomberg’s K statistics (Blomberg et al. 2003). We cross-validated those two indices because their estimation might be affected by branch lengths uncertainties in time-calibrated phylogenies (Molina-Venegas and Rodríguez 2017). To account for tree topology uncertainties, we calculated both indices using 1,000 trees from the posterior distribution and we calculated their mean values and 95% confidence limits. Pagel’s λ ranges from 0 to 1, with increasing values indicating stronger phylogenetic signal. K ranges from 0 (no phylogenetic signal) to infinity, with K=1 suggesting that the trait has evolved according to the Brownian motion model of evolution and K values >1 suggesting that closely related species are more similar than expected under the Brownian motion model (Blomberg et al. 2003).

The evolution of the climatic traits and the elevation was visualised by ancestral reconstructions of mean values of the four bioclimatic variables, along the species-level time-calibrated tree using the fastAnc() function in the R package Phytools. Elevation can be used as another proxy of species climatic and habitat requirements (Yu et al. 2016), thus, we also carried out niche evolution analyses using the mean elevation of species occurrences as another trait.

### Climatic niche

We described and compared climatic niches of individual species using the R package ecospat v. 3.2 (di Cola et al. 2017). First, we delimited background climate to describe the full range of conditions over the entire range of the genus *Erebia*, as recommended by Broennimann et al., (2012). The background climatic data were extracted from a rectangle delimited by the minimum and maximum longitude and latitude of the occurrence points. We extracted values of the four bioclimatic variables from >100,000 points in a regular grid covering the rectangle. The background dataset is used to infer null model niches built by random sampling from the background area (di Cola et al., 2017). Second, we added data on the four selected bioclimatic variables for the occurrence points of all species to the background dataset. Then, PCA was used to transform *n* correlated variables into two uncorrelated linear combinations (principal components) of the original variables (di Cola et al., 2017). These values describe the climatic niches of individual species. We further calculated niche width for each species and niche overlap for species pairs following Broennimann et al. (2012), see below for more details.

### Niche width in the European and the Asian clade

The range of climatic conditions inhabited by individual species is described as the climatic niche width. First, we calculated the total climatic niche width based on the PCA of the 4 bioclimatic variables. Species with ≤ 5 distinct geo-referenced occurrence records and associated climate data were excluded; thus, we kept 68 species in our climatic niche width analyses. We calculated the relative area of the polygon described by the species-specific occurrence points in the twodimensional representation of its niche estimated by PCA (Broennimann et al. 2012). Second, we calculated niche width separately for each of the four bioclimatic variables and for the elevation. The niche width for individual variables was approximated using the 0.05 and 0.95 quantiles of the variable to limit the influence of outliers on our niche width estimates. Annual precipitation (bio12) was log_10_-transformed prior to niche width calculation.

We used generalised linear models (GLM) with Gaussian error distribution to compare the climatic niche width between species from the European and the Asian clades. To account for the potential confounding effect of the number of records on species’ niche width estimates, we also included the number of occurrence points (log-transformed) and membership in the Asian or the European clade for each species as additional explanatory variables in the GLMs. Next, we tested whether species’ niche width depends on the position of their niche along the gradients of individual climate variables and along the altitudinal gradient, i.e., whether species occurring in more extreme conditions are more specialised. To test this hypothesis, we fitted a generalised additive model (GAM, R package mgcv v. 1.8-38, Wood 2017) with Gaussian error distribution and a smooth function (cubic spline) describing the changes of species’ niche width along the gradient of individual climate variables or elevation. We fitted a separate model for each variable.

### Climatic niche diversification

Divergence of the climatic niche between sister species, particularly when they have overlapping distribution ranges (broadly defined as sympatry here), might represent a signature of climate-driven differentiation expected in the adaptive radiation scenario. To test this hypothesis, we calculated the mean overlap of climatic niches for all sister vs. non-sister species pairs and for sympatric vs. allopatric (i.e. no spatial overlap of their ranges) sister species pairs. For each species pair, we calculated the overlap of the total climatic niche and, separately, the niche overlaps for each of the four climate variables and the species’ elevational ranges. We followed the approach of Broennimann et al. (2012). We extracted PCA scores of occurrence locations for individual species and calculated the occurrence densities in a grid across the first two PCA axes using the resolution of 100×100 pixels (Broennimann et al., 2012; di Cola et al., 2017). We kept the 68 species with > 5 occurrence points in our dataset for analysis. Then, we calculated the total climatic niche overlap of all species pairs using the Schoener’s D index following Bandeira et al. (2021). The resulting pairwise matrix contained climatic niche overlap values for all species pairs (Supplementary material, Table S5). We separately calculated the overlap of each of the four bioclimatic variables (Supplementary material, Tables S6-S9) and the overlaps of the elevational ranges (Supplementary material, Table S10) of individual species pairs. We calculated the overlap of each variable for all species pairs again using the Schoener’s D index.

To compare the niche overlap values (i.e., the estimated Schoener’s D) between sister species pairs and non-sister species pairs, we carried out two-tailed Mantel tests with 9,999 permutations using the ecodist v. 2.0.7 package for R (Goslee and Urban 2007, Goslee 2010). We compared the pairwise climatic niche overlap matrix (Supplementary materials, Table S5-S10) against a matrix containing the value of 1 for sister species pairs and the value of 0 for all non-sister species pairs (Supplementary material, Table S11). Furthermore, we compared the values of the niche overlaps between sympatric and allopatric sister species pairs (Supplementary material, Table S12) using generalised linear models (GLM) with Gaussian error distribution. This analysis included 16 sister species pairs after the exclusion of species with an insufficient number of occurrence points (≤ 5 distinct locations).

## RESULTS

### Phylogenetic tree and biogeography

The age of origin of extant *Erebia* lineages was estimated at 22.4 ± 4.60 Ma. The two main clades, the European clade and the Asian clade, diverged during the Early to Middle Miocene, 16 ± 4 Ma (Figure 1; Supplementary material, Figure S1 for the normal distribution and Figure S2 for the uniform distribution of the calibration points). The European clade, including the *tyndarus* group, had a crown age of 15 ± 4 Ma. Most extant lineages diversified during the Late Miocene and the Pliocene (5 – 10 Ma).

**Figure 1.**
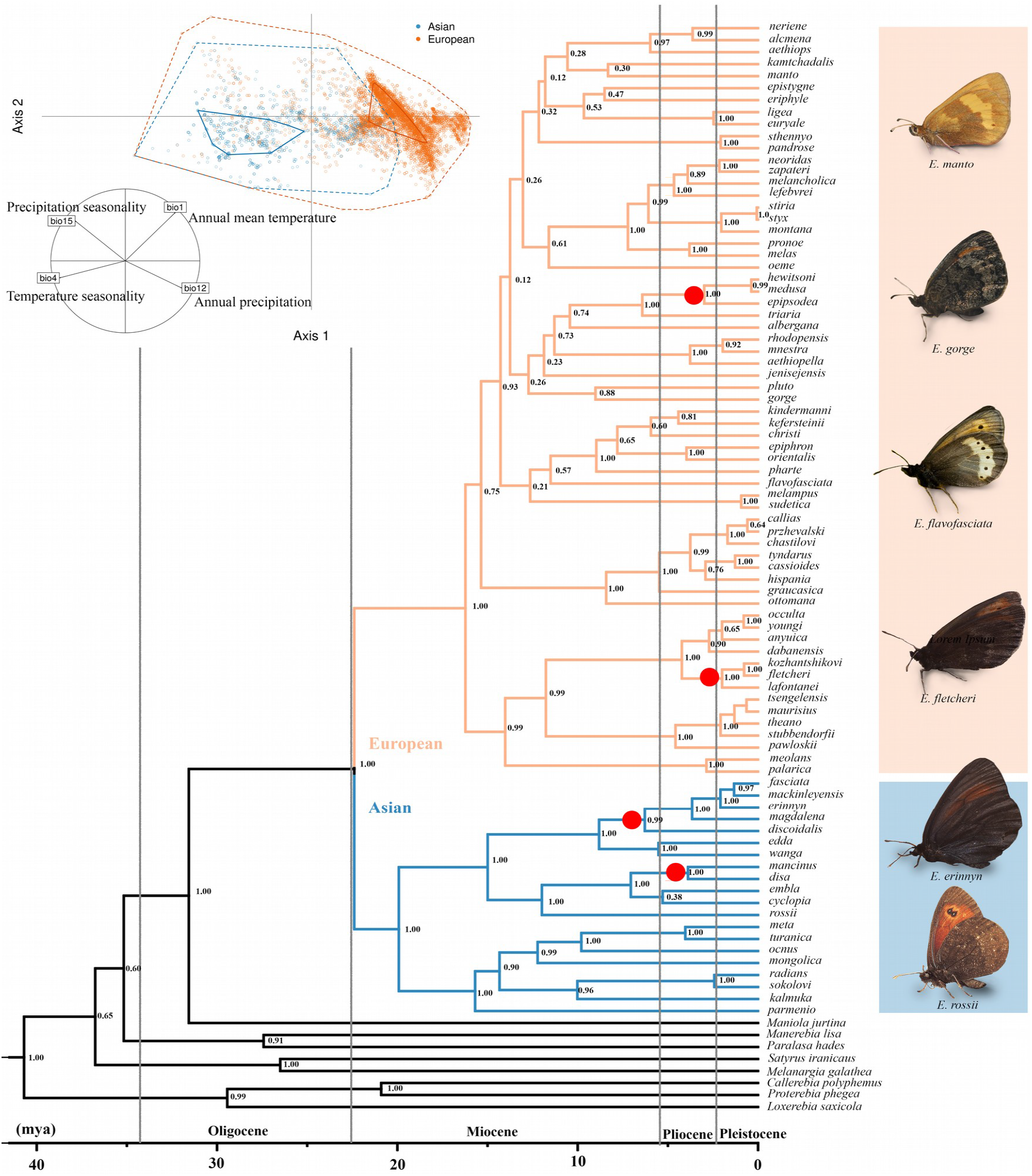
Phylogenetic relationships and divergence time estimates for the butterfly genus *Erebia*. The European clade diversified mainly in Europe (orange), the Asian clade in Asia (blue), although the geographic distribution of several species is not restricted to a single region (see Supplementary material, Figure S3). The full biogeographic reconstruction is shown in the Supplementary material, Figures S3-S5. Climatic PCA displays the distribution-based climatic data for all species of the European and the Asian clades. The polygons show the full extent of the conditions occupied by each clade (dotted lines) and the core 50% of the climatic niche (solid lines). The inset shows the correlation of individual bioclimatic variables with the first and second PCA axes. The bioclimatic variables (bio1, bio4, bio12 and bio15) displayed significant phylogenetic signal (Table 3). The European clade inhabits warmer, more humid, and less seasonal climate compared to the Asian clade.

The best-fit biogeographical model was the BayArea-like+J (J = 0.05) followed by DEC+J (J = 0.03) (ΔAICc = 6.4) and DEC models (ΔAICc = 12.7) (Supplementary material, Table S13). The estimated ancestral areas were consistent across all tested models, except of the ancestral area of the genus at the root of the phylogeny. The best-fit model (BayArea-like+J) estimated that the ancestral area was the Eastern Palaearctic (Supplementary material, Figure S3), whereas the DEC+J and the DEC models (Supplementary material, Figures S4 and S5) estimated a widespread distribution in the Eastern and Western Palaearctic for the crown *Erebia*. Members of the European clade expanded their distributions toward North America at least 5 times independently since 7 Ma (Figure S3). The representatives of the Asian clade further diversified within the region, and they expanded toward North America twice, at about 3 Ma.

### Evolution of the climatic traits

The climatic PCA analysis of distribution-based climatic data (Figure 1) suggested that members of the European clade inhabit warmer and more humid climates with lower seasonality, whereas members of the Asian clade inhabit places with colder, dryer, and more seasonal conditions. The comparison of mean values and ranges of the four bioclimatic variables and the elevation of the distribution-based data for the European and the Asian clade confirmed this pattern (Table 1).

**Table 1.**
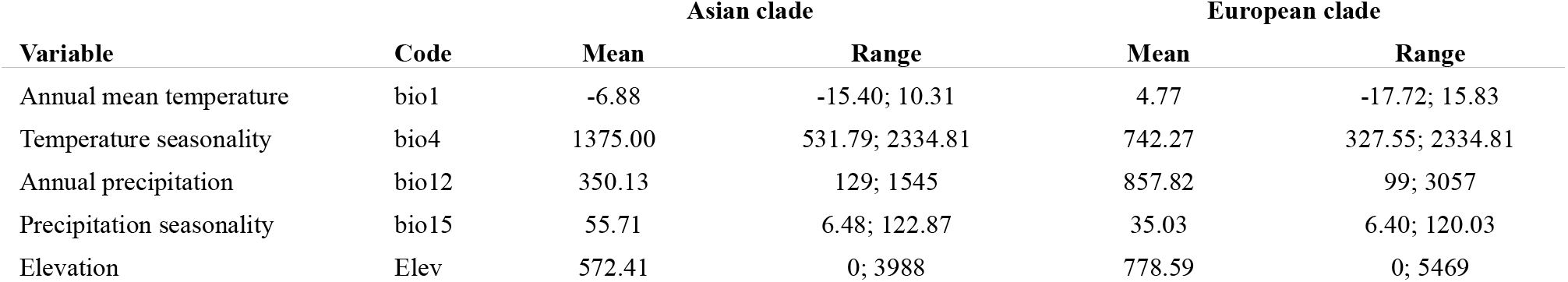
Overview of the climatic conditions and elevation at sites where species from the Asian clade and the European clade of the butterfly genus *Erebia* were recorded. Mean values and range of the four bioclimatic variables and the elevation associated with individual occurrence records of all species, divided into the two clades, are summarised.

The evolution of climatic traits was better fitted by the Ornstein-Uhlenbeck (OU) and Brownian motion (BM) models than by the Early Burst model. The annual mean temperature (bio1), annual precipitation (bio12) and precipitation seasonality (bio15) likely evolved toward local optima, given the better fit of the OU model (Table 2), though the deviation from Brownian motion, described by the parameter α, was low (α < 0.2). The evolution of both temperature seasonality (bio4) and elevation were described comparably well by both the OU, with little deviation from Brownian motion, and the Brownian motion models (Δ AICc < 2). We used the OU models for reconstructions of all climatic traits, as these models had the best fit to our data.

**Table 2.** Comparison of the fit of three models of trait evolution, Brownian Motion (BM), Ornstein-Uhlenbeck (OU), and Early Burst (EB) for the five WorldClim variables across the phylogeny of the genus *Erebia*. The rate parameter (σ^2^) describes how fast the trait values change through time. θ is mean value of a trait, α describes the strength of attraction toward the mean value in the OU model, and a is the rate change parameter in the EB model. Models with the best fit (ΔAICc < 2) are highlighted in bold.

| Variable | Model | $\sigma^2$ | $\theta$ | $\alpha$ | A | logL | AICc | $\Delta AICc$ |
| --- | --- | --- | --- | --- | --- | --- | --- | --- |
| bio1 | BM | 2.20 | -1.32 | – | – | -232.31 | 468.77 | 5.18 |
|  | **OU** | **3.21** | **-0.89** | **0.060** | – | **-228.64** | **463.58** | **0.00** |
|  | EB | 2.20 | -1.32 | – | -0.000001 | -232.31 | 470.92 | 7.34 |
| bio4 | **BM** | **6083.2** | **1069.7** | – | – | **-561.16** | **1126.47** | **0.00** |
|  | **OU** | **6933.0** | **1060.7** | **0.017** | – | **-560.84** | **1127.98** | **1.51** |
|  | EB | 6083.3 | 1069.7 | – | -0.000001 | -561.16 | 1128.62 | 2.15 |
| bio12 | BM | 21159.8 | 642.5 | – | – | -612.89 | 1229.93 | 22.57 |
|  | **OU** | **37149.4** | **735.5** | **0.123** | – | **-600.53** | **1207.36** | **0.00** |
|  | EB | 21160.3 | 642.5 | – | -0.000001 | -612.89 | 1232.09 | 24.72 |
| bio15 | BM | 75.8 | 53.1 | – | – | -379.19 | 762.53 | 24.28 |
|  | **OU** | **149.4** | **46.3** | **0.150** | – | **-365.97** | **738.25** | **0.00** |
|  | EB | 75.8 | 53.1 | – | -0.000001 | -379.19 | 764.68 | 26.43 |
| Elevation | **BM** | **54548.0** | **1525.5** |  |  | **-652.19** | **1308.53** | **1.94** |
|  | **OU** | **73081.7** | **1547.7** | **0.043** |  | **-650.14** | **1306.59** | **0.00** |
|  | EB | 54549.1 | 1525.5 |  | -0.000001 | -652.19 | 1310.69 | 4.09 |

The evolution of the climatic traits of the genus *Erebia* held strong phylogenetic signal for the λ and the K indices, when analysed for the consensus tree (Table 3). The narrow confidence limits of λ and K calculated based on 1000 randomly selected trees from the posterior distribution of phylogenetic trees suggested a consistent phylogenetic signal across estimated tree topologies. When analysed across posterior distribution of phylogenetic trees, the mean annual temperature (bio1), temperature seasonality (bio4), and elevation held strong phylogenetic signal in both λ and K, whereas annual precipitation (bio12) and precipitation seasonality (bio15) held phylogenetic signal only recovered by the λ index. The reconstruction of the ancestral climatic trait values (Figure 2, panels C – G) is consistent with the evolution of the climatic niches toward different local optima in the two main clades (Table 2). As indicated by the PCA analysis (see above), the ancestral reconstruction of climatic traits shows that, compared to their common ancestor, the members of the Asian clade evolved climatic niches characterised by colder (Figure 2C), dryer (Figure 2F), and more seasonal climate (Figure 2D, G), while the European clade shifted toward warmer, more humid, and less seasonal climate. The ancestral reconstruction of the elevational niche showed that related species occur in similar elevations (Figure 2E), with low elevations colonised in the cold Arctic areas and in warm (Figure 2C) areas of the temperate zone.

**Figure 2.**
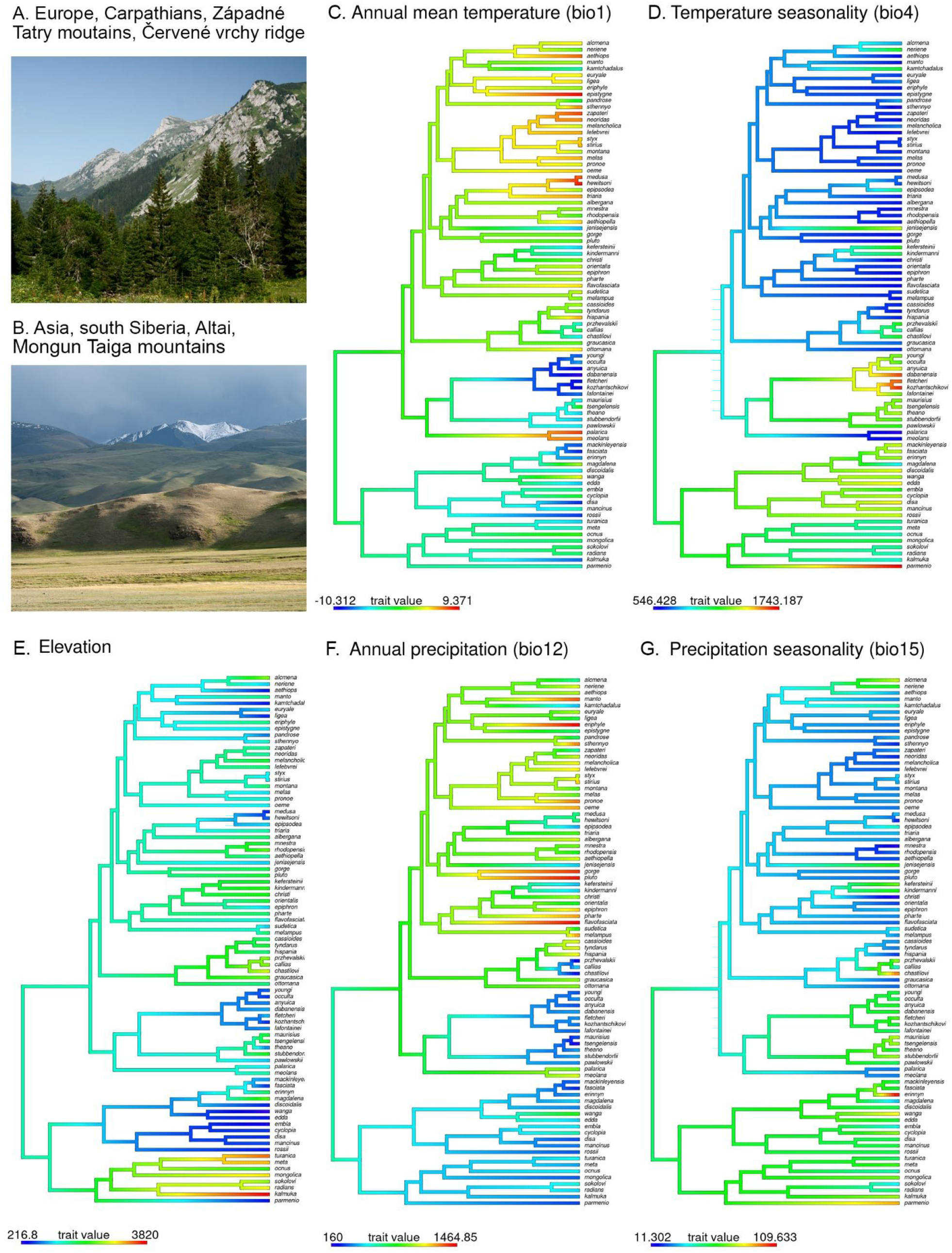
Illustrative figures of European and Asian mountains (panels A, B) and ancestral reconstruction of climatic traits (panels C-G) for species-specific average values of WorldClim variables. The ancestral reconstructions are based on OU model of evolution in the butterfly genus *Erebia*.

**Table 3.**
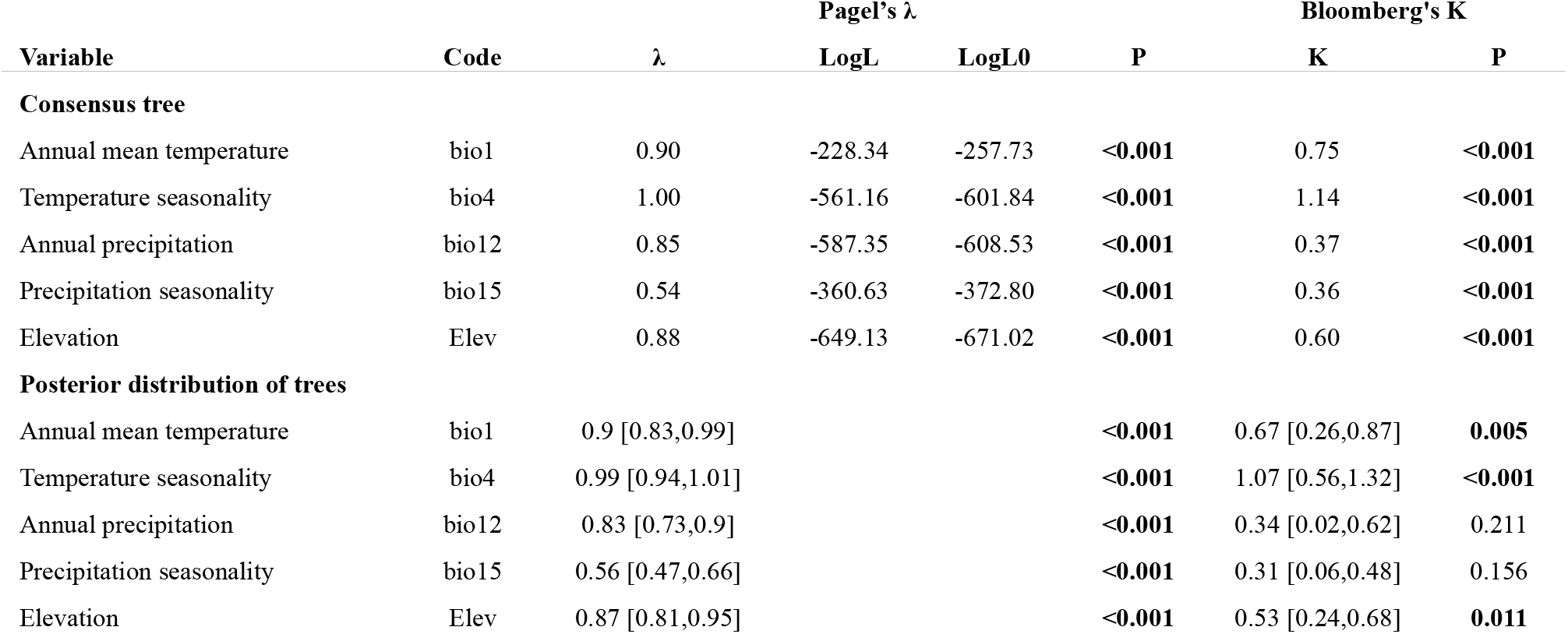
Tests of phylogenetic signal in climatic traits and elevation in the butterfly genus *Erebia* for the consensus tree and for 1,000 trees randomly drawn from the posterior distribution (mean value and 95% confidence interval). The climatic and elevation traits were computed as average values of distribution-based climate data (WorldClim variables bio1, bio4, bio12, bio15, and elevation). LogL = log likelihood of the model with the estimated value of the Pagel’s λ, LogL0 = log likelihood of a model with Pagel’s λ=0. The presence of a significant phylogenetic signal is highlighted in bold.

### Climatic niche width

Comparison of niche width between species in the Asian clade and the European clade showed that the species of the European clade had narrower thermal niches based on annual mean temperature and temperature seasonality (Table 4). The width of the precipitation niche, the total niche inferred from the climatic PCA, and the elevational range were not significantly different between the two clades (Table 4). In addition, the GAM models testing for changes of species niche width across climatic gradients (Figure 3) indicated that the climatic niche was narrower, thus, species were more specialised, under extreme environmental conditions, particularly strongly in the case of mean temperature.

**Figure 3.**
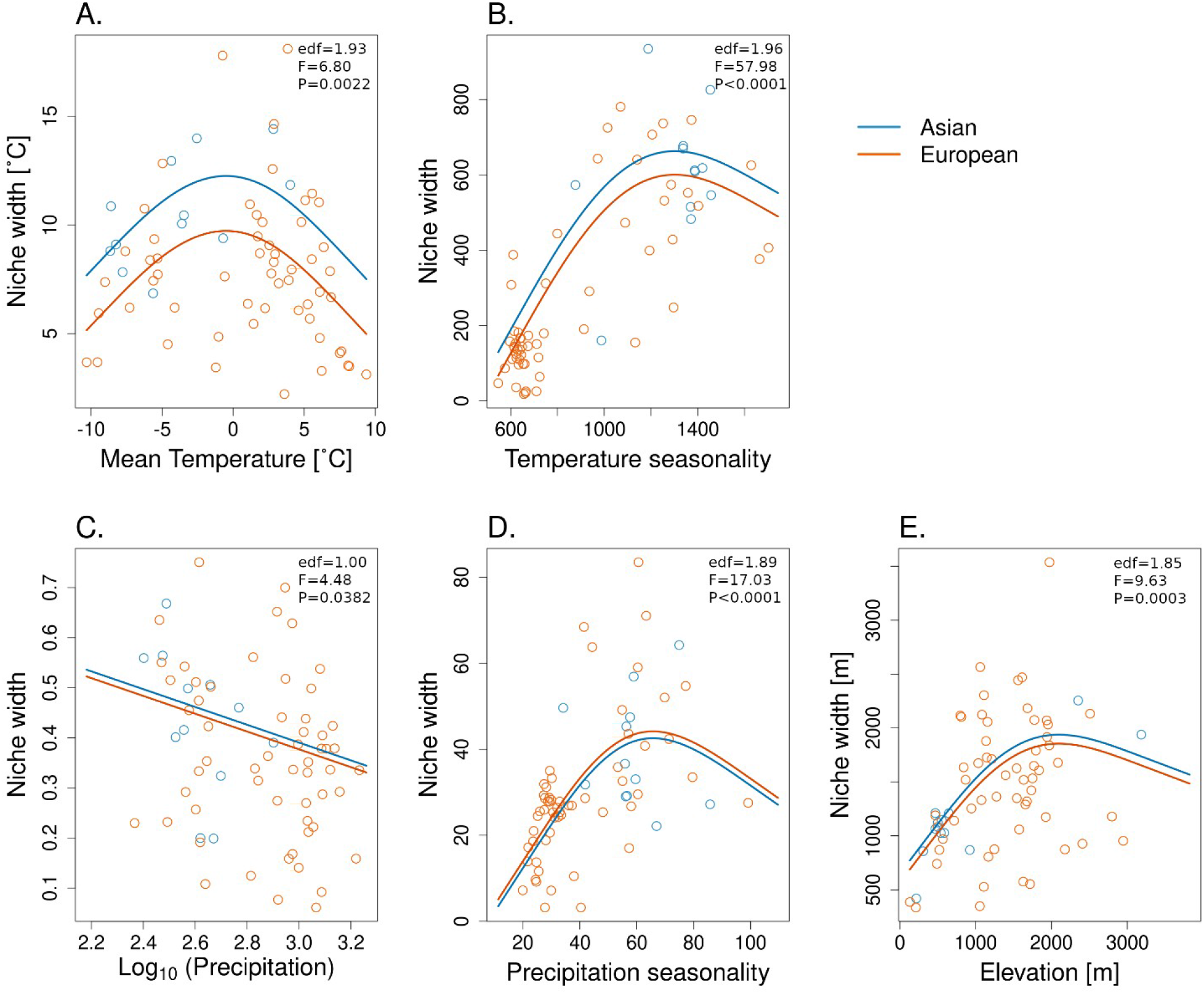
The relationships between climatic and elevation niche widths and niche position along the gradient of conditions occupied by the butterfly genus *Erebia*. The points correspond to the niche width (y-axis) and average value of a climate variable or elevation for individual species (x-axis). The best fit of generalised additive models (GAM) is shown by lines (separately for species of the European and the Asian clade). Statistical significance of the model fit is also shown. edf denotes the estimated degrees of freedom, which indicates the complexity of the non-linear relationship (edf = 1 corresponds to a linear relationship).

**Table 4.**
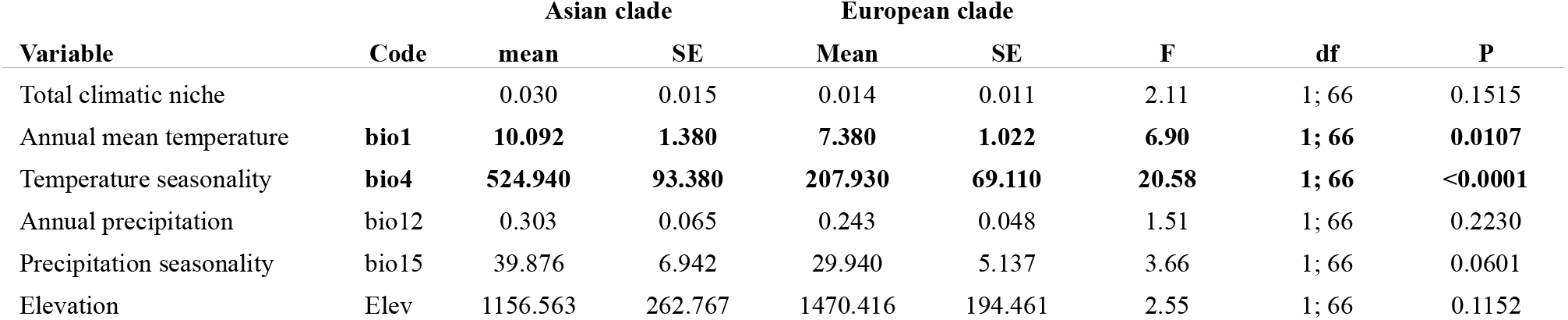
Comparison of climatic and elevation niche widths of species from the Asian clade and the European clade of the butterfly genus *Erebia*. Niche width was calculated for the total climatic niche based on PCA of four climate variables, for the four variables separately, and for the elevation. Mean niche width and standard error are shown for the two clades. Annual precipitation was log_10_-transformed prior to the calculation of niche width. Results of generalised linear models (GLM) with Gaussian error distribution testing the difference of the climatic niche width between the European and Asian clades are shown. The number of occurrence points (log-transformed) with climate data for each species was also included in the models to account for the potential confounding effect on species’ niche width. Significant differences in niche width between the Asian and the European clade are highlighted in bold.

| Variable | Code | Asian clade |  | European clade |  | F | df | P |
| --- | --- | --- | --- | --- | --- | --- | --- | --- |
|  |  | mean | SE | Mean | SE |  |  |  |
| Total climatic niche |  | 0.030 | 0.015 | 0.014 | 0.011 | 2.11 | 1; 66 | 0.1515 |
| Annual mean temperature | **bio1** | **10.092** | **1.380** | **7.380** | **1.022** | **6.90** | **1; 66** | **0.0107** |
| Temperature seasonality | **bio4** | **524.940** | **93.380** | **207.930** | **69.110** | **20.58** | **1; 66** | **<0.0001** |
| Annual precipitation | bio12 | 0.303 | 0.065 | 0.243 | 0.048 | 1.51 | 1; 66 | 0.2230 |
| Precipitation seasonality | bio15 | 39.876 | 6.942 | 29.940 | 5.137 | 3.66 | 1; 66 | 0.0601 |
| Elevation | Elev | 1156.563 | 262.767 | 1470.416 | 194.461 | 2.55 | 1; 66 | 0.1152 |

### Climatic niche diversification

The mean overlap of climatic niches between sister species was significantly higher than in nonsister species pairs for all climatic variables (Mantel test, P < 0.05 in all cases, Table 5). However, we have not detected any significant differences in the niche overlap between sympatric and allopatric sister species pairs, although we note that the number of species we could include in the analysis (16 sister species pairs) was rather low (GLM, P > 0.05 in all cases, Supplementary material, Table S14).

**Table 5.** Comparison of mean overlap of the climatic niche, indicated by Schoeners’ D values, between sister species pairs and non-sister species pairs of the butterfly genus *Erebia*. The niche overlap was calculated for the total climatic niche based on the PCA of four climate variables, for each variable separately, and for elevation. Annual precipitation was log_10_-transformed. Two-tailed Mantel test was used to compare the values of niche overlap between sister and non-sister species pairs. The Schoeners’ D values were calculated only for species with > 5 distribution points; in total the analyses included 68 species.

| Variable | Code | Sister species pairs |  | Non-sister species pairs |  | Mantel r | P |
| --- | --- | --- | --- | --- | --- | --- | --- |
|  |  | mean | SD | Mean | SD |  |  |
| Total climatic niche |  | 0.309 | 0.232 | 0.123 | 0.170 | 0.089 | **0.0001** |
| Annual mean temperature | bio1 | 0.465 | 0.258 | 0.278 | 0.249 | 0.062 | **0.0016** |
| Temperature seasonality | bio4 | 0.471 | 0.214 | 0.194 | 0.245 | 0.093 | **0.0001** |
| Annual precipitation | bio12 | 0.458 | 0.235 | 0.226 | 0.233 | 0.082 | **0.0003** |
| Precipitation seasonality | bio15 | 0.428 | 0.307 | 0.237 | 0.239 | 0.065 | **0.0012** |
| Elevation | Elev | 0.470 | 0.222 | 0.292 | 0.237 | 0.062 | **0.0010** |

## DISCUSSION

The extant species diversity of mountain butterflies in the genus *Erebia* was likely shaped by climatic niche conservatism within local optima (Figure 2, Table 2) and adaptation to local climates in specific lineages (Table 1). The conservatism within the two main *Erebia* clades might have resulted in repeated cycles of range expansion and isolation linked to past climatic fluctuations (Schmitt et al. 2016). Climatic niche specialisation might have further contributed to the occupancy of regions with climatic extremes at the margins of genus’ distribution across the Holarctic (Figure 3), in terms of annual mean temperature, seasonality of temperature and precipitation, and elevation. Species in the European clade are characterized by narrower range of temperature and its seasonality compared to members of the Asian clade (Table 4). Also, the precipitation niche was narrower in the high-precipitation (mainly European) and broader in low-precipitation (mainly Asian) parts of the precipitation gradient (Figure 3, panel C).

Spatial sorting (Comerford and Egan 2022) might have preceded climatic niche specialization in the European clade followed by species differentiation (Vamosi et al. 2014). On the other hand, high species richness and competition for resources in zones of secondary contact might have driven niche specialization by adaptive radiation to decrease the level of competition (Vázquez and Stevens 2004). The adaptive radiation scenario in the genus *Erebia* has been suggested in the light of gradual slow-down of diversification rate within the genus (Peña et al. 2015). However, we did not find support for rapid diversification of the climatic niche early in the evolution of *Erebia* (represented by the Early Burst model), which is not consistent with the adaptive radiation scenario. We did find support for phylogenetic signal and the absence of climatic niche diversification in sister species pairs, including those with sympatric ranges, which suggests conservatism in climatic traits. However, as niche is multidimensional and our study includes only several dimensions of the niche space, adaptive divergence could appear in other traits, such as habitat use.

### Climatic niche conservatism as the primary driver of speciation

The major and rapid radiation of *Erebia* in the Middle Miocene (Figure 1) was concurrent with the uplift of the main mountain massifs in the Palearctic, such as the Qinghai-Tibetan Plateau (Valdiya 2016), European Alps (Ager 1975, Botsyun et al. 2020), and the Caucasus (Hrivniak et al. 2020). The newly arising mountain areas have likely promoted a burst of speciation in *Erebia* (Peña et al. 2015), similarly to other Palearctic mountains butterflies (Su et al. 2020), mayflies (Hrivniak et al. 2020), and plants (Hörandl and Emadzade 2012, Folk et al. 2019). Although the climate was warmer during the Miocene and the Pliocene than during the Pleistocene (Botsyun et al. 2020), the isolated mountain ridges could have represented islands of grassland habitats necessary for *Erebia*. Indeed, grassland habitats spread in Europe since the Late Miocene, about 8.5 Ma (Feurdean and Vasiliev 2019). More recent climatic changes during the Pleistocene have further driven diversification among populations at the intraspecific level (Louy et al., 2014a, Louy, et al., 2014b, Wendt et al., 2022), which maintained the rich species diversity established since the Miocene.

The climatic niche evolution seems to be directed toward different local optima in Europe and Asia as suggested by OU models of trait diversification (Table 2), but the deviation from the Brownian motion model of trait evolution was small. The niche conservatism (Table 3) in both the European (wet and less seasonal climatic niche) and the Asian clade (dryer and more seasonal climatic niche), might have driven allopatric speciation thanks to dispersal and isolation of populations during past climatic fluctuations (Louy, et al., 2014b). Similarly, climatic niche conservatism has been linked to allopatric speciation in temperate groups (Wiens and Graham 2005), such as insular bees (Dorey et al. 2020), mountain rodents (Wan et al. 2018), and micropterigid moths in the Japanese Archipelago (Imada et al. 2011). In line with this, the climatic niche similarity found between sister species (Table 5) reinforces the hypothesis that *Erebia* species likely diversified in allopatry (or parapatry) (Schmitt et al. 2016) rather than by rapid climatic niche evolution (lability) (Pitteloud et al. 2017) which would have resulted in substantial speciation rates in sympatry. The degree of conservatism quantified here favours the hypothesis that the conserved climatic niche coupled with climate change could be a driver of speciation in these mountain butterflies. However, even within the European clade, there is a significant variation in the species niche, with several sister species pairs where one species occupies colder conditions than the other (e.g. sister species pairs *E. ligea* and *E. euryale*, or *E. pandrose* and *E. sthennyo;* see Fig. 2). Hence, it seems that the diversification of *Erebia* was driven by complex events linked to differentiation across distinct climatic niches and specific habitats (e.g. grasslands, gravel, woodland clearings, Varga, 2014).

### Niche specialisation contributed to species diversification

Specialisation to climatic extremes in *Erebia* is evidenced by the hump-shaped dependence of niche width on climatic gradients (Figure 3). We detected higher niche specialisation under extreme values of annual mean temperature, temperature- and precipitation-seasonality, and elevation. The interpretation of this finding is that taxa occupying areas with extreme climatic conditions tend to have narrow niches. This pattern has been found in amphibians (Bonetti and Wiens 2014), but rarely documented in temperate insects. It indicates that the occupancy of both climatic margins, warm lowlands and cold mountain summits, demands physiological specialisation, which might have prevented the colonization of such areas by taxa having broad climatic niches. Indeed, we previously observed distinct thermoregulatory adaptations in European *Erebia* species in alpine and lowland habitats (Kleckova et al. 2014, Kleckova & Klecka 2016). Niche specialisation has been considered an evolutionary dead-end (Colles et al. 2009, Devictor et al. 2010) due to the tendency of specialists to inhabit smaller geographical ranges, which may enhance extinction rates. Adaptations to extreme conditions found in populations inhabiting Palearctic mountain summits have also been detected in *Parnassius* butterflies (Su et al. 2020), whose species diversification in mountains was likely regulated by ecological limits (Condamine 2018). The linear decrease of precipitation niche width with increasing precipitation (Figure 3, panel C) could perhaps reflect diversification by divergence in other traits, such as habitat use, in areas with higher precipitation. Such a pattern in climatic niche linked to precipitation has been also found in salamanders across a wide range of latitudes (Bonetti and Wiens 2014).

### The sources of European diversity

The higher species diversity of the genus *Erebia* in Europe than in Asia is an exceptional pattern of biodiversity distribution in the Holarctic (Sanmartin et al. 2001). In the species rich European clade, we detected generally narrower thermal niches than in the Asian clade (Figure 3, panel A). We argue that such pattern may relate to the west-east seasonality gradient, with lower seasonality in Europe than in Asia (Table 1). The higher variability of climatic conditions connected with climatic seasonality is assumed to cause general widening of climatic niches, such that temperate taxa are expected to have broader climatic niches than tropical ones (MacArthur 1972). In line with this expectation, the temperature seasonality niche width positively correlates with temperature seasonality, with only a few species occurring in the most seasonal conditions deviating from this pattern (Figure 3, panel B).

The disparate patterns in temperature niche width found in the European and Asian clades can be explained by several ecological and evolutionary mechanisms. For example, the narrower temperature niches of European lineages might have resulted from a diversity-dependent radiation event (Peña et al. 2015), suggesting that competition for resources or other ecological processes have driven niche specialisation. In the case of polyphagous butterflies, such as the genus *Erebia*, potential competition for resources is not limited to larval host plants but can be also related to the availability of nectar, microhabitat use (Kleckova et al., 2014), and competition for mates. Alternatively, the narrower niches can be seen as a “by-product” of allopatric speciation across and within mountain ranges, without any signs of adaptive or ecological processes. Under this scenario, species diversification might be explained by genetic drift in molecular (Lucek, 2018) or reproductive traits (Schat et al. 2022), and random evolution of the climatic niche.

### Study limitations and future perspectives

The macroevolutionary and macroecological patterns that we detected are robust. However, to further understand the processes leading to speciation in *Erebia*, we still need more detailed information about the ecology (e.g., habitat use or potential host plant specialisation), physiology, morphology, and genome evolution of *Erebia* species. First, the climatic data was interpreted at a macroscale which might have limited the detection of ecological divergence in locally sympatric species. We estimated climatic niches on a relatively rough scale of 1 km^2^, which does not capture fine-scale microclimatic differentiation (Kirchheimer et al. 2016). It is thus expected that regionally sympatric species might also display finer niche partitioning and differentiation of their realized climatic and microhabitat niches (Lorkovic 1958, Kleckova et al. 2014), which might have undetected by our study (Supplementary material, Table S8). In addition, different *Erebia* species seem to have diversified in their adult phenologies, which might have further created niche partitioning (van Dam et al. 2019). Further, environmental conditions not included in our study, such as winter temperature minima or low snow cover (Konvicka et al. 2021), may affect more vulnerable stages of the developmental cycle, such as overwintering larvae (Vrba et al. 2017), and affect species distributions and the observed climatic niches. Last, the evolution of the climatic niche is difficult to separate from biogeographical histories as both processes are interconnected (Coelho et al. 2019, Comerford and Egan 2022).

Our estimates of the timing of diversification events in *Erebia* (see also Peña et al. 2015) suggest that the origin of the majority of European species preceded the Pleistocene (during Miocene and Pliocene), but Schmitt et al. (2016) considered a Pleistocene origin of the rich diversity of *Erebia*.

Time calibration used in our study is based on fossil-based secondary calibrations and a phylogenetic tree inferred from mitochondrial and nuclear genes (Chazot et al. 2021). The alternative time scenario is based on substitution rates of two mitochondrial genes, ND5 and CoxII (Albre et al., 2008, Schmitt et al. 2016). However, the evolution and substitution rates of mitochondrial genes can be affected by presence of Wolbachia endosymbionts and the selective sweeps of associated mitochondria (Suchackova et al. 2021). For example, we inferred a crown age of 8.5 (+-3) Ma for the *tyndarus* species group, but Albre et al. (2008) suggested the Pleistocene origin (<2.8 Ma). However, our hypothesis is supported also by a study comparing genomes of secondary contact zones of European butterflies (Ebdon et al., 2021). Although estimates of the timing of diversification events provide useful context for the interpretation of our results, our main conclusions do not depend on the time scenario.

The origin of alpine endemic insects has complex drivers (Queiroz et al. 2022), including allopatric speciation (Schoville and Roderick, 2010, Schat et al. 2022, Todisco et al. 2010), diversity dependent radiation (Condamine 2018), or interactions with endosymbionts (Lucek et al. 2021, Weng et al. 2021). We fill in new information into the mosaic describing diversification drivers of alpine insects in relation to the role of the evolution of their climatic niches. We demonstrated, first, the evolution of climatic niches towards local optima in response to geographical shifts (see also Kleckova et al. 2015) and second, climatic niche specialisation along the gradient of temperature suggesting a trade-off in adaptation to low- and high-temperature limits. Further, the climatic specialisation detected in the European clade could reflect divergence in other traits such as habitat use. Finally, we conclude that bridging the gap between studies of population- and species-level differentiation (Capblancq et al. 2019) will be a fruitful avenue for future research to resolve the sources of the extraordinary diversity of the genus *Erebia*, as well as other alpine insects.

## ACKNOWLEDGEMENTS

Funding was provided by the Czech Science Foundation (GAČR grant: GJ20-18566Y), Computational resources were supplied by the project “e-Infrastruktura CZ” (e-INFRA CZ LM2018140) supported by the Ministry of Education, Youth and Sports of the Czech Republic. We thank Jiri Skala for providing additional *Erebia* specimens and Alena Suchackova-Bartonova for help in a laboratory. We are grateful for constructive comments provided by three anonymous reviewers and the editors.

