## Supplementary material for "Climatic niche conservatism and ecological diversification in the Holarctic cold-dwelling butterfly genus *Erebia*": Kleckova_Erebia_Supplementary materials_revised.pdf

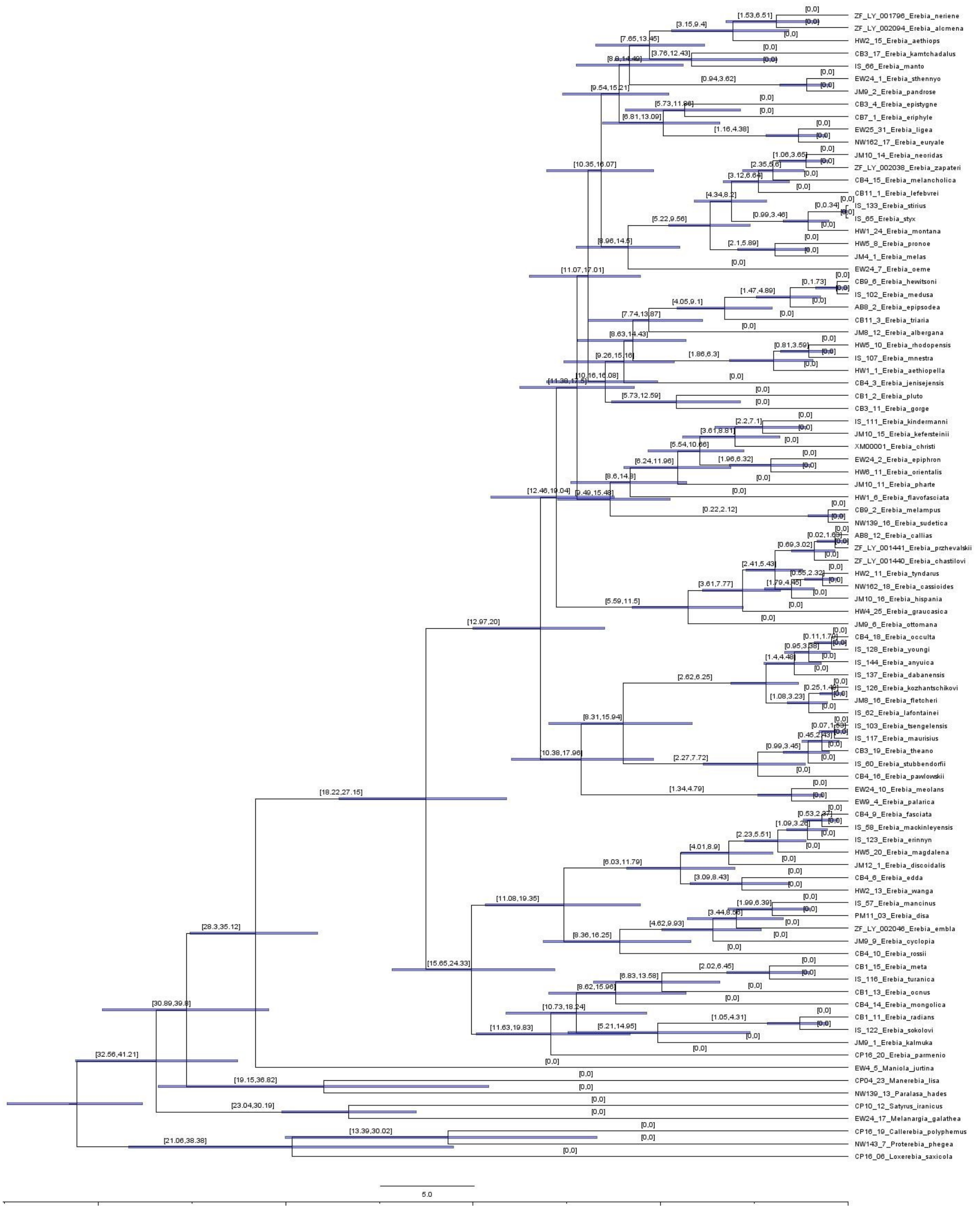


**Figure S1.** Phylogenetic relationships and divergence time estimates of the genus *Erebia* inferred by BEAST2 with *normal* distribution used as a prior for the age of the calibrated nodes. Node bars and the numbers above the nodes show the 95% highest posterior density interval to indicate the uncertainty of the estimates.

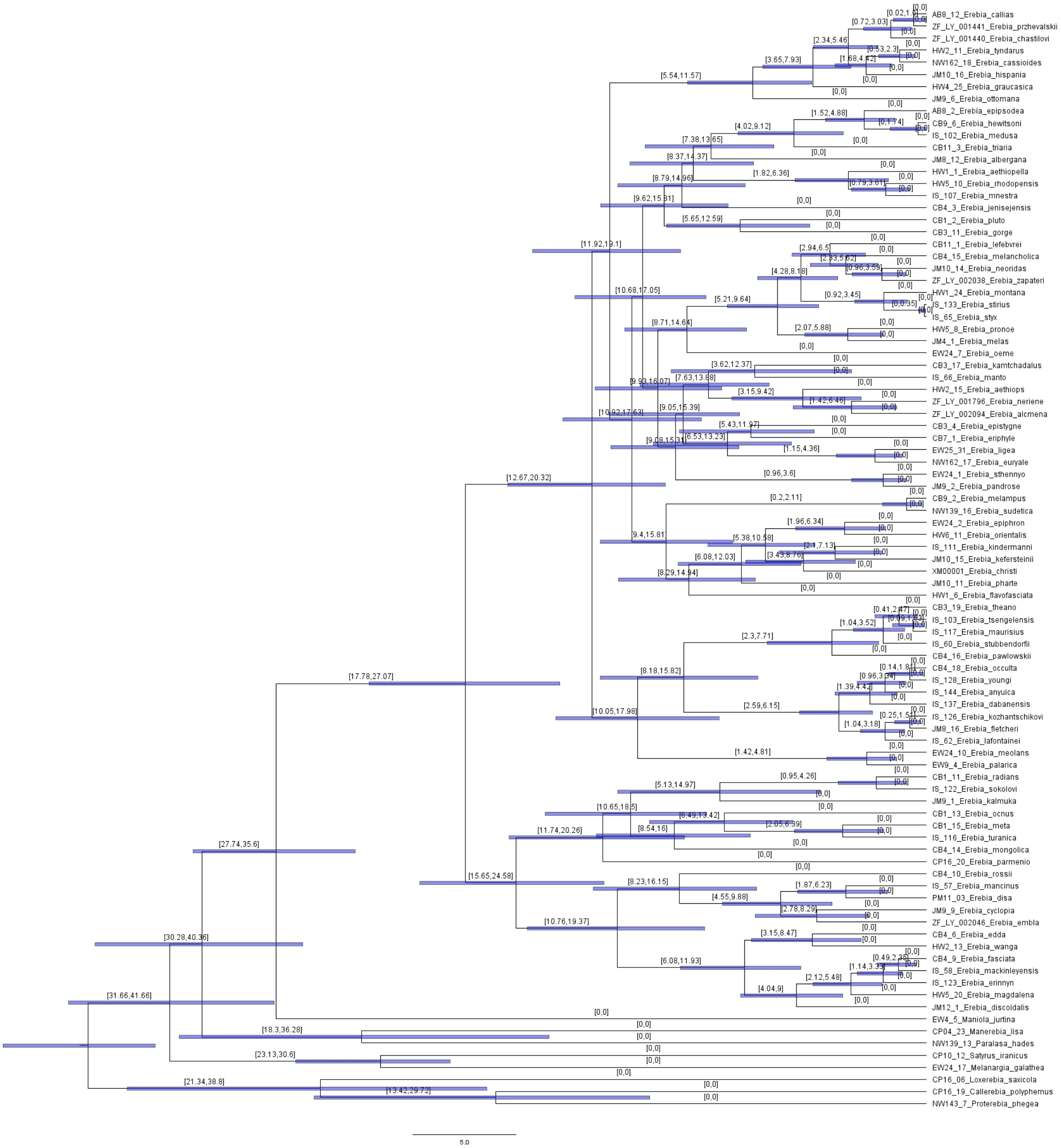


**Figure S2.** Phylogenetic relationships and divergence time estimates of the genus *Erebia* inferred by BEAST2 with *uniform* distribution used as a prior for the age of calibrated nodes. Node bars and the numbers above the nodes show the 95% highest posterior density interval to indicate the uncertainty of the estimates.

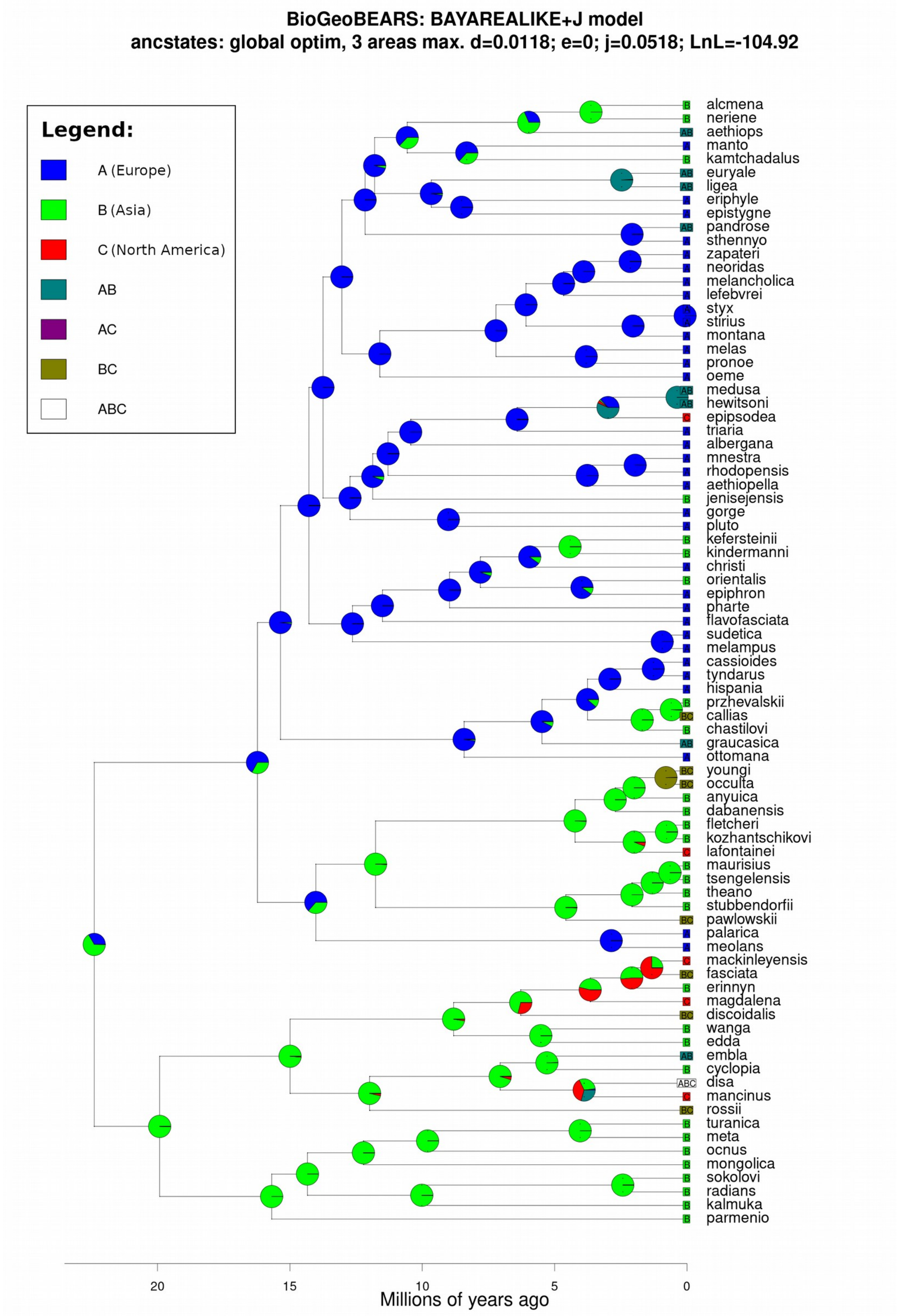


**Figure S3.** Biogeographic reconstruction of the butterfly genus *Erebia* analysed by the BioGeoBEARS software based on the best-fit model BAYAREALIKE+J. The species and ancestral nodes with predicted occurrence in Asia (B) are in green, in Europe (A) in blue and in North America (C) are in red. The comparison of tested models is presented in Table S13.

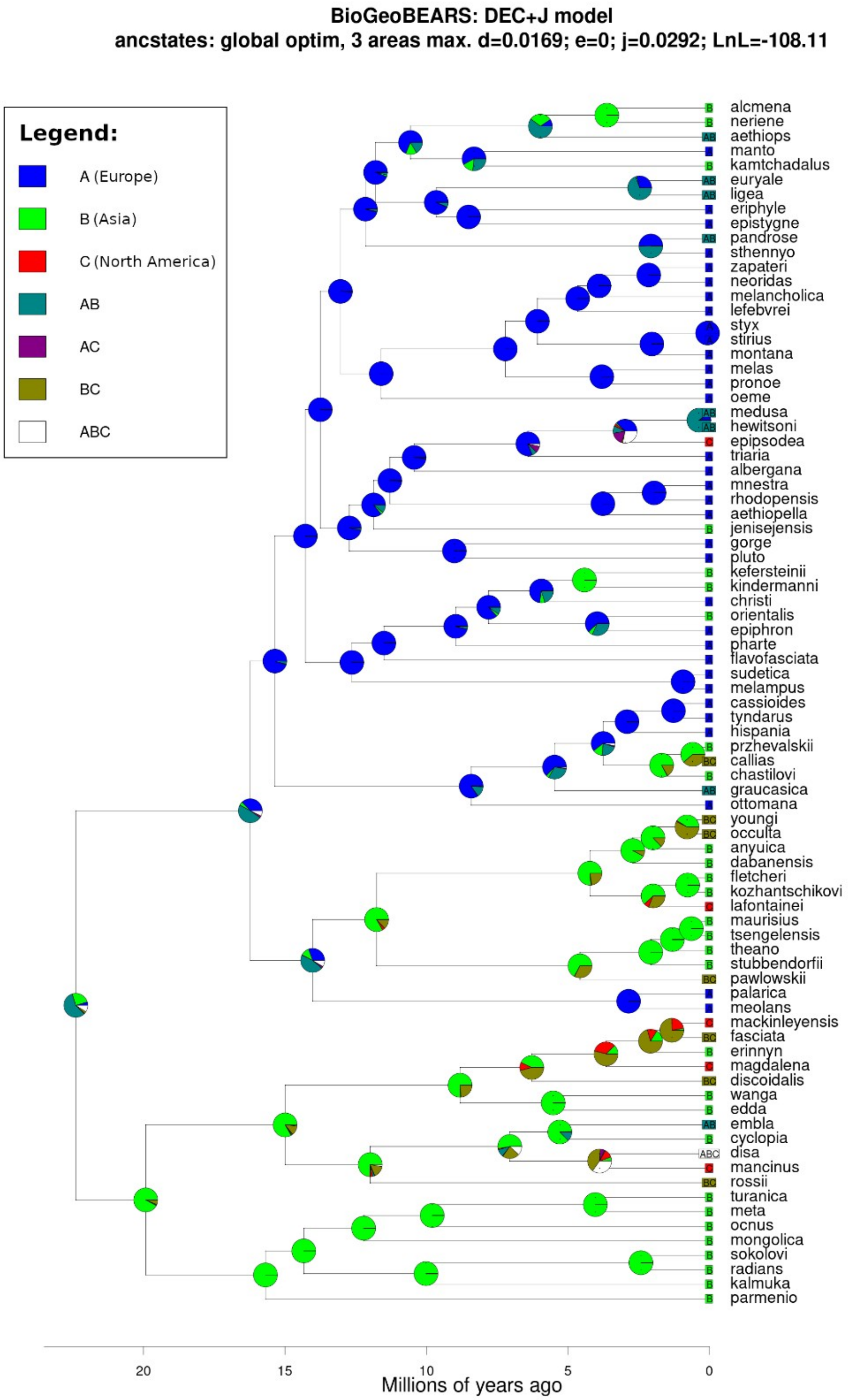


**Figure S4.** Biogeographic reconstruction of the butterfly genus *Erebia* analysed by the BioGeoBEARS software based on the model DEC+J. The species and ancestral nodes with predicted occurrence in Asia (B) are in green, in Europe (A) in blue and in North America (C) are in red. The comparison of tested models is presented in Table S13.

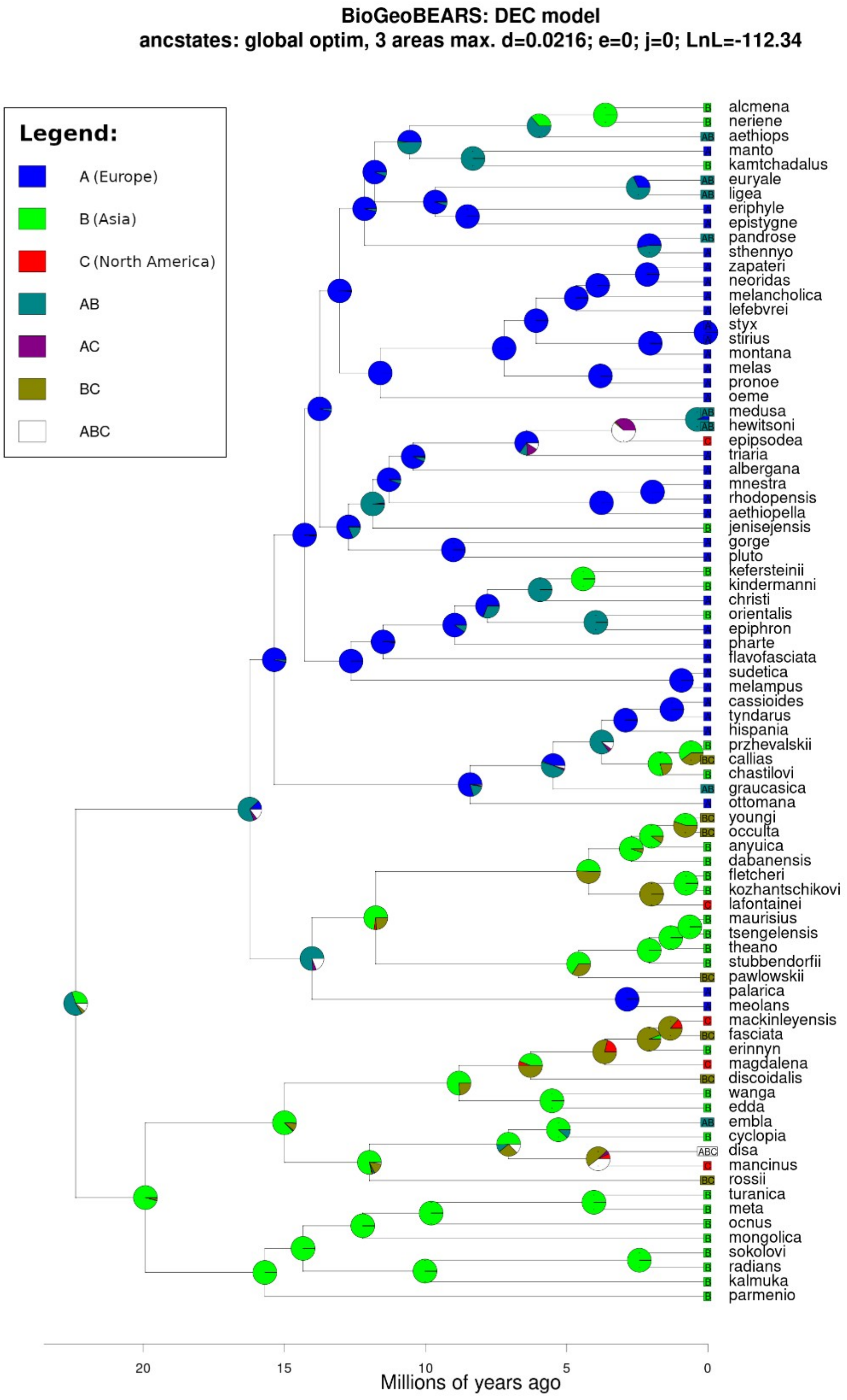


**Figure S5.** Biogeographic reconstruction of the butterfly genus *Erebia* analysed by the BioGeoBEARS software based on the model DEC. The species and ancestral nodes with predicted occurrence in Asia (B) are in green, in Europe (A) in blue and in North America (C) are in red. The comparison of tested models is presented in Table S13.

**Table S1.** Specimen list used in the study including the specimen code, country of origin, and GenBank accession numbers for four genes used for phylogenetic reconstruction (COI – in two parts, GAPDH, RpS5, and wingless).

| Code | Subtribe | Genus | Species | Country | COI-begin | COI-end | GAPDH | RpS5 | wingless |
|---|---|---|---|---|---|---|---|---|---|
| NW139-13 | Ypthimina | Paralasa | hades | Tajikistan | GQ357215 | - | GQ357453 | GQ357582 | GQ357348 |
| EW24-17 | Melanargiina | Melanargia | galathea | France | DQ338843 | DQ338843 | EU528398 | EU528444 | DQ338706 |
| EW4-5 | Maniolina | Maniola | jurtina | Spain | AY090214 | AY090214 | EU141481 | EU141376 | KR139116 |
| NW143-7 | Maniolina | Proterebia | afra | Greece | GQ357221 | GQ357221 | GQ357460 | GQ357590 | GQ357353 |
| CP04-23 | Pronophilina | Manerebia | lisa | Peru | GQ357233 | GQ357233 | GQ357480 | GQ357609 | GQ357366 |
| CP16-06 | Ypthimina | Loxerebia | saxicola | China | GQ357214 | GQ357214 | GQ357451 | GQ357580 | GQ357347 |
| CP16-19 | Ypthimina | Callerebia | polyphemus | China | GQ357212 | GQ357212 | GQ357449 | GQ357578 | GQ357345 |
| CP10-12 | Satyrina | Satyrus | iranicus | Iran | KR138743 | KR138743 | GQ357508 | GQ357634 | DQ338740 |
| HW1-1 | Erebiina | Erebia | aethiopella | Italy | KR138756 | - | KR139008 | - | KR139118 |
| HW2-15 | Erebiina | Erebia | aethiops | Switzerland | KR138768 | KR138768 | KR139018 | KR138903 | - |
| JM8-12 | Erebiina | Erebia | albergana | Austria | KR138816 | - | KR139056 | KR138946 | KR231856 |
| ZF-LY-002094 | Erebiina | Erebia | alcmena | China | X | - | - | - | - |
| IS-144 | Erebiina | Erebia | anyuica | Russia | MW318418 | MW318418 | MW317722 | MW317457 | MW318708 |
| AB8-12 | Erebiina | Erebia | callias | USA | KR138709 | - | - | - | KR139077 |
| NW162-18 | Erebiina | Erebia | cassioides | France | KR138836 | KR138836 | KR139071 | KR138966 | KR231857 |
| ZF-LY-001440 | Erebiina | Erebia | chastilovi | Mongolia | KR138842 | - | - | - | KR139166 |
| XM00001 | Erebiina | Erebia | christi | Italy | MN829479.1 | - | - | - | - |
| JM9-9 | Erebiina | Erebia | cyclopia | Russia | KR138829 | KR138829 | - | KR138959 | KR139161 |
| IS-137 | Erebiina | Erebia | dabanensis | Russia | MW318419 | MW318419 | KR139034 | KR138919 | KR139148 |
| PM11-03 | Erebiina | Erebia | disa | Canada | KR138841 | KR138841 | KR139075 | KR138971 | KR139165 |
| JM12-1 | Erebiina | Erebia | discoidalis | Canada | KR138807 | KR138807 | KR139048 | KR138937 | KR231860 |
| CB4-6 | Erebiina | Erebia | edda | Russia | KR138736 | - | KR138998 | KR138879 | KR139106 |
| ZF-LY-002046 | Erebiina | Erebia | embla | Russia | X | - | - | - | X |
| EW24-2 | Erebiina | Erebia | epiphron | France | KR138751 | KR138751 | KR139006 | KR138888 | KR139113 |
| AB8-2 | Erebiina | Erebia | epipsodea | USA | KR138712 | - | - | - | KR139080 |
| CB3-4 | Erebiina | Erebia | epistygne | Spain | KR138727 | - | KR138991 | KR138872 | KR139099 |
| IS-123 | Erebiina | Erebia | erinnyn | Russia | KR138789 | KR138789 | KR139031 | KR138917 | KR139145 |
| CB7-1 | Erebiina | Erebia | eriphyle | Austria | KR138738 | - | KR139000 | KR138881 | KR139108 |
| NW162-17 | Erebiina | Erebia | euryale | France | KR138835 | KR138835 | KR139070 | KR138965 | KR231863 |
| CB4-9 | Erebiina | Erebia | fasciata | Canada | KR138737 | - | KR138999 | KR138880 | KR139107 |
| HW1-6 | Erebiina | Erebia | flavofasciata | Italy | KR138764 | - | - | KR138899 | KR139126 |
| JM8-16 | Erebiina | Erebia | fletcheri | Russia | KR138819 | - | KR139059 | KR138949 | KR139158 |
| CB3-11 | Erebiina | Erebia | gorge | Switzerland | KR138720 | KR138720 | KR138986 | KR138867 | KR139094 |
| HW4-25 | Erebiina | Erebia | graucasica | Armenia | KR138770 | KR138770 | KR139020 | KR138905 | KR139131 |
| CB9-6 | Erebiina | Erebia | hewitsoni | Turkey | X | - | - | - | - |
| JM10-16 | Erebiina | Erebia | hispania | Spain | KR138803 | KR138803 | KR139045 | KR138932 | KR231866 |
| CB4-3 | Erebiina | Erebia | jenisejensis | Russia | KR138735 | - | KR138997 | KR138878 | KR139105 |
| JM9-1 | Erebiina | Erebia | kalmuka | Kyrgyzstan | KR138821 | - | - | KR138951 | - |
| JM10-15 | Erebiina | Erebia | kefersteini | Russia | KR138802 | KR138802 | - | KR138931 | KR231867 |
| IS-111 | Erebiina | Erebia | kindermanni | Russia | KR138787 | KR138787 | - | - | KR139141 |
| IS-126 | Erebiina | Erebia | kozhantshikovi | Russia | KR138790 | KR138790 | KR139032 | - | KR139146 |
| IS-62 | Erebiina | Erebia | lafontainei | USA | KR138795 | KR138795 | KR139038 | KR138923 | KR139152 |
| CB11-1 | Erebiina | Erebia | lefebvrei | Spain | KR138713 | KR138713 | KR138976 | KR138857 | KR139084 |
| CB3-17 | Erebiina | Erebia | kamtschadalis | Russia | KR138723 | - | KR138989 | KR138870 | KR139097 |
| EW25-31 | Erebiina | Erebia | ligea | Sweden | KR138753 | KR138753 | - | KR138890 | KR139115 |
| IS-58 | Erebiina | Erebia | mackinleyensis | Canada | KR138793 | KR138793 | KR139036 | KR138921 | KR139150 |
| HW5-20 | Erebiina | Erebia | magdalena | USA | KR138777 | KR138777 | KR139023 | KR138909 | KR139135 |
| IS-57 | Erebiina | Erebia | mancinus | Canada | KR138792 | KR138792 | KR139035 | KR138920 | KR139149 |
| IS-66 | Erebiina | Erebia | manto | Italy | - | KR138845 | KR139041 | KR138926 | KR139155 |
| IS-117 | Erebiina | Erebia | maurisius | Russia | MW318424 | MW318424 | - | KR138915 | KR139143 |
| IS-102 | Erebiina | Erebia | medusa | Russia | KR138784 | KR138784 | - | KR138914 | KR139139 |
| CB9-2 | Erebiina | Erebia | melampus | Austria | KR138739 | KR138739 | KR139001 | KR138882 | KR139109 |
| CB4-15 | Erebiina | Erebia | melancholica | Russia | KR138732 | - | KR138996 | KR138877 | KR139104 |
| JM4-1 | Erebiina | Erebia | melas | Bulgaria | KR138812 | KR138812 | KR139052 | KR138942 | KR231873 |
| EW24-10 | Erebiina | Erebia | meolans | France | KR138749 | KR138749 | - | KR138886 | KR139112 |
| CB1-15 | Erebiina | Erebia | meta | Kazakhstan | KR138716 | KR138716 | KR138979 | KR138860 | KR139087 |
| IS-107 | Erebiina | Erebia | mnestra | Italy | KR138786 | KR138786 | - | - | - |
| CB4-14 | Erebiina | Erebia | mongolica | Kyrgyzstan | KR138731 | KR138731 | KR138995 | KR138876 | KR139103 |

| | | | | | | | | | |
|---|---|---|---|---|---|---|---|---|---|
| HW1-24 | Erebiina | Erebia | montana | Italy | KR138762 | - | KR139013 | KR138896 | KR139123 |
| JM10-14 | Erebiina | Erebia | neoridas | Italy | KR138801 | KR138801 | KR139044 | KR138930 | KR139156 |
| ZF-LY-001796 | Erebiina | Erebia | neriene | Russia | X | - | - | - | X |
| CB4-18 | Erebiina | Erebia | occulta | Canada | KR138734 | - | - | - | - |
| CB1-13 | Erebiina | Erebia | ocnus | Kazakhstan | KR138714 | KR138714 | KR138977 | KR138858 | KR139085 |
| EW24-7 | Erebiina | Erebia | oeme | France | DQ338780 | DQ338780 | EU141479 | EU141375 | DQ338640 |
| HW6-11 | Erebiina | Erebia | orientalis | Bulgaria | KR138782 | KR138782 | KR139027 | - | - |
| JM9-6 | Erebiina | Erebia | ottomana | Macedonia | KR138826 | KR138826 | KR139064 | KR138956 | KR231878 |
| EW9-4 | Erebiina | Erebia | palarica | Spain | KR138755 | KR138755 | GQ357422 | GQ357551 | KR139117 |
| JM9-2 | Erebiina | Erebia | pandrose | Russia | KR138823 | KR138823 | - | KR138953 | KR231880 |
| CP16-20 | Erebiina | Erebia | parmenio | Russia | KR138746 | - | KR139004 | - | - |
| CB4-16 | Erebiina | Erebia | pawloskii | Canada | KR138733 | - | - | - | - |
| JM10-11 | Erebiina | Erebia | pharte | Switzerland | KR138799 | - | - | KR138928 | - |
| CB1-2 | Erebiina | Erebia | pluto | Italy | KX040618 | KR138846 | KR138981 | KR138862 | KR139089 |
| HW5-8 | Erebiina | Erebia | pronoe | Bulgaria | KR138780 | KR138780 | KR139026 | KR138912 | KR139138 |
| ZF-LY-001441 | Erebiina | Erebia | przhevalskii | Mongolia | KR138843 | - | - | - | KR139167 |
| CB1-11 | Erebiina | Erebia | radians | Kazakhstan | - | KR138849 | KR138975 | KR138856 | KR139083 |
| HW5-10 | Erebiina | Erebia | rhodopensis | Bulgaria | KR138773 | KR138773 | - | - | KR139133 |
| CB4-10 | Erebiina | Erebia | rossii | Canada | KR138729 | - | KR138993 | KR138874 | KR139101 |
| IS-122 | Erebiina | Erebia | sokolovi | Kyrgyzstan | - | KR138850 | - | KR138916 | KR139144 |
| EW24-1 | Erebiina | Erebia | sthennyo | France | KR138748 | KR138748 | KR139005 | KR138885 | DQ338641 |
| IS-133 | Erebiina | Erebia | stiria | Slovenia | MN752712 | MN752712 | - | - | - |
| IS-60 | Erebiina | Erebia | stubbendorfii | Russia | KR138794 | KR138794 | KR139037 | KR138922 | KR139151 |
| IS-65 | Erebiina | Erebia | styx | Austria | KR138797 | KR138797 | KR139040 | KR138925 | KR139154 |
| NW139-16 | Erebiina | Erebia | sudetica | Czechia | KR138830 | - | - | KR138960 | - |
| CB3-19 | Erebiina | Erebia | theano | Kazakhstan | KR138724 | - | - | - | - |
| CB11-3 | Erebiina | Erebia | triaria | Spain | KR138715 | - | KR138978 | KR138859 | KR139086 |
| IS-103 | Erebiina | Erebia | tsengelensis | Mongolia | KR138785 | KR138785 | KR139029 | - | KR139140 |
| IS-116 | Erebiina | Erebia | turanica | Kyrgyzstan | KR138788 | KR138788 | KR139030 | - | KR139142 |
| HW2-11 | Erebiina | Erebia | tyndarus | Switzerland | KR138766 | KR138766 | KR139016 | KR138901 | KR139128 |
| HW2-13 | Erebiina | Erebia | wanga | Russia | KR138767 | KR138767 | KR139017 | KR138902 | KR139129 |
| IS-128 | Erebiina | Erebia | youngi | Canada | KR138791 | KR138791 | KR139033 | KR138918 | KR139147 |
| ZF-LY-002038 | Erebiina | Erebia | zapateri | Spain | X | - | - | - | X |

**Table S2.** Alignment partitions for mitochondrial (COI*,* position 1-1475) and nuclear (GAPDH, 1476-2166; RpS5*,* 2168-2783 and *wingless,* 2783-3195) genes specified by the software PartitionFinder 2. For mitochondrial genes we detected low effective sample size (ESS) values. Thus, we reduced the complexity of models on substitution models for HKY for the mitochondrial subset1 and HKY+G+I in the mitochondrial subsets 2 and 3.

| Gene | Character set |
|---|---|
| *Mitochondrial* | 1Subset = 1-1475\3; |
| *Mitochondrial* | 2Subset = 2-1475\3; |
| *Mitochondrial* | 3Subset = 3-1475\3; |
| *Nuclear* | 4Subset = 2167-2783\3 1476-2166\3; |
| *Nuclear* | 5Subset = 2168-2783\3 1477-2166\3 2786-3195\3 2785-3195\3; |
| *Nuclear* | 6Subset = 1478-2166\3 2169-2783\3; |
| *Nuclear* | 7Subset = 2784-3195\3; |

| Subset | Best Model | sites | Partition names | |
|---|---|---|---|---|
| 1 | GTR+G+X | 492 | COIcodon1 | reduced to HKY |
| 2 | TRN+I+G+X | 492 | COIcodon2 | reduced to HKY+G+I |
| 3 | TRN+I+G+X | 491 | COIcodon3 | reduced to HKY+G+I |
| 4 | HKY+G+X | 437 | Rp5Scodon1, GAPDHcodon1 | |
| 5 | TRN+G+X | 710 | Rp5Scodon2, GAPDHcodon2, winglesscodon3, winglesscodon2 | |
| 6 | JC+I | 435 | GAPDHcodon3, Rp5Scodon3 | |
| 7 | HKY+G+X | 138 | winglesscodon1 | |

**Table S3.** Biogeographic distribution of the genus *Erebia* in the three main biogeographic regions, A-Europe, B-Asia, C-North America, in the PHYLIP format, used for analyses of biogeography in the R package BioGeoBEARS v1.1.2 (Matzke 2013).

| 83 | 3 (A B C) |
|---|---|
| *aethiopella* | 100 |
| *aethiops* | 110 |
| *albergana* | 100 |
| *alcmena* | 010 |
| *anyuica* | 010 |
| *callias* | 011 |
| *cassioides* | 100 |
| *chastilovi* | 010 |
| *christi* | 100 |
| *cyclopia* | 010 |
| *dabanensis* | 010 |
| *disa* | 111 |
| *discoidalis* | 011 |
| *edda* | 010 |
| *embla* | 110 |
| *epiphron* | 100 |
| *epipsodea* | 001 |
| *epistygne* | 100 |
| *erinnyn* | 010 |
| *eriphyle* | 100 |
| *euryale* | 110 |
| *fasciata* | 011 |
| *flavofasciata* | 100 |
| *fletcheri* | 010 |
| *gorge* | 100 |
| *graucasica* | 110 |
| *hewitsoni* | 110 |
| *hispania* | 100 |
| *jenisejensis* | 010 |
| *kalmuka* | 010 |
| *kamtschadalis* | 010 |
| *kefersteinii* | 010 |
| *kindermanni* | 010 |
| *kozhantshikovi* | 010 |
| *lafontainei* | 001 |
| *lefebvrei* | 100 |
| *ligea* | 110 |
| *mackinleyensis* | 001 |
| *magdalena* | 001 |
| *mancinus* | 001 |
| *manto* | 100 |
| *maurisius* | 010 |
| *medusa* | 110 |
| *melampus* | 100 |
| *melancholica* | 100 |

| | |
|---|---|
| *melas* | 100 |
| *meolans* | 100 |
| *meta* | 010 |
| *mnestra* | 100 |
| *mongolica* | 010 |
| *montana* | 100 |
| *neoridas* | 100 |
| *neriene* | 010 |
| *occulta* | 011 |
| *ocnus* | 010 |
| *oeme* | 100 |
| *orientalis* | 010 |
| *ottomana* | 100 |
| *palarica* | 100 |
| *pandrose* | 110 |
| *parmenio* | 010 |
| *pawloskii* | 011 |
| *pharte* | 100 |
| *pluto* | 100 |
| *pronoe* | 100 |
| *przhevalskii* | 010 |
| *radians* | 010 |
| *rhodopensis* | 100 |
| *rossii* | 011 |
| *sokolovi* | 010 |
| *sthennyo* | 100 |
| *stiria* | 100 |
| *stubbendorfii* | 010 |
| *styx* | 100 |
| *sudetica* | 100 |
| *theano* | 010 |
| *triaria* | 100 |
| *tsengelensis* | 010 |
| *turanica* | 010 |
| *tyndarus* | 100 |
| *wanga* | 010 |
| *youngi* | 011 |
| *zapateri* | 100 |

**Table S4.** Occurrence records of the all species of the butterfly genus *Erebia* used in this study. Because of taxonomic uncertainties, we included only North American occurrence records of *E. pawloskii,* as its records from Asia are probably frequently confused with records of related *E. stubbendorfii*. *Provided as a separate file.*

**Table S5.** The matrix of total climatic niche overlap values for all *Erebia* species pairs estimated by the Schoener's D index. *Provided as a separate file.*

**Table S6.** The matrix of bio1 niche overlap values for all *Erebia* species pairs estimated by the Schoener's D index. *Provided as a separate file.*

**Table S7.** The matrix of bio4 niche overlap values for all *Erebia* species pairs estimated by the Schoener's D index. *Provided as a separate file.*

**Table S8.** The matrix of bio12 niche overlap values for all *Erebia* species pairs estimated by the Schoener's D index. *Provided as a separate file.*

**Table S9.** The matrix of bio15 niche overlap values for all *Erebia* species pairs estimated by the Schoener's D index. *Provided as a separate file.*

**Table S10**. The matrix of the elevation niche overlap values for all *Erebia* species pairs estimated by the Schoener's D index. *Provided as a separate file.*

**Table S11.** Comparison of the niche overlap matrix with *Erebia* species x species matrix which contained the value of 1 for sister species pairs and the value of 0 for all non-sister species pairs. *Provided as a separate file.*

**Table S12.** The list of all sister species pairs of *Erebia* based on the topology of the consensusn phylogenetic tree (Figure S1). Niche overlap is indicated by Schoeners' D values. The Schoeners' D values were calculated for pairs of species where both species had > 5 distribution points. Niche overlap was calculated for the total climatic niche based on the PCA of four climate variables, for each variable separately, and for the elevation. Annual precipitation was $\log_{10}$-transformed. We also classified the species pairs as either sympatric or allopatric.

| Sister species pair | | D(climate) | D(bio1) | D(bio4) | D(bio12) | D(bio15) | D(elevation) | Distribution |
|---|---|---|---|---|---|---|---|---|
| *neriene* | *alcmena* | *NA* | *NA* | *NA* | *NA* | *NA* | *NA* | allopatry |
| *kamtschadalis* | *manto* | *NA* | *NA* | *NA* | *NA* | *NA* | *NA* | allopatry |
| *epistygne* | *eriphyle* | 0.052 | 0.013 | 0.152 | 0.051 | 0.754 | 0.322 | allopatry |
| *ligea* | *euryale* | 0.512 | 0.822 | 0.468 | 0.595 | 0.687 | 0.443 | sympatry |
| *sthennyo* | *pandrose* | *NA* | *NA* | *NA* | *NA* | *NA* | *NA* | allopatry |
| *neoridas* | *zapateri* | 0.000 | 0.178 | 0.218 | 0.299 | 0.017 | 0.582 | allopatry |
| *stiria* | *styx* | 0.140 | 0.506 | 0.745 | 0.801 | 0.174 | 0.454 | sympatry |
| *pronoe* | *melas* | 0.414 | 0.655 | 0.537 | 0.366 | 0.535 | 0.788 | sympatry |
| *hewitsoni* | *medusa* | *NA* | *NA* | *NA* | *NA* | *NA* | *NA* | sympatry |
| *rhodopensis* | *mnestra* | *NA* | *NA* | *NA* | *NA* | *NA* | *NA* | allopatry |
| *pluto* | *gorge* | 0.692 | 0.838 | 0.728 | 0.726 | 0.792 | 0.829 | sympatry |
| *kindermanni* | *kefersteinii* | 0.401 | 0.385 | 0.634 | 0.255 | 0.614 | 0.298 | sympatry |
| *epiphron* | *orientalis* | *NA* | *NA* | *NA* | *NA* | *NA* | *NA* | allopatry |
| *melampus* | *sudetica* | 0.010 | 0.210 | 0.129 | 0.036 | 0.090 | 0.087 | allopatry |
| *przhevalskii* | *callias* | *NA* | *NA* | *NA* | *NA* | *NA* | *NA* | sympatry |
| *tyndarus* | *cassioides* | 0.638 | 0.783 | 0.771 | 0.751 | 0.690 | 0.743 | allopatry |
| *occulta* | *youngi* | 0.536 | 0.688 | 0.681 | 0.617 | 0.667 | 0.801 | sympatry |
| *kozhantshikovi* | *fletcheri* | 0.294 | 0.518 | 0.500 | 0.337 | 0.144 | 0.327 | sympatry |
| *tsengelensis* | *maurisius* | *NA* | *NA* | *NA* | *NA* | *NA* | *NA* | allopatry |
| *meolans* | *palarica* | 0.161 | 0.535 | 0.204 | 0.550 | 0.102 | 0.418 | sympatry |
| *fasciata* | *mackinleyensis* | 0.521 | 0.475 | 0.556 | 0.532 | 0.784 | 0.269 | sympatry |
| *edda* | *wanga* | 0.030 | 0.099 | 0.427 | 0.293 | 0.028 | 0.389 | sympatry |
| *mancinus* | *disa* | 0.360 | 0.254 | 0.444 | 0.481 | 0.622 | 0.506 | sympatry |
| *cyclopia* | *embla* | 0.182 | 0.480 | 0.342 | 0.637 | 0.150 | 0.265 | sympatry |
| *meta* | *turanica* | *NA* | *NA* | *NA* | *NA* | *NA* | *NA* | allopatry |
| *radians* | *sokolovi* | *NA* | *NA* | *NA* | *NA* | *NA* | *NA* | sympatry |

**Table S13**. Comparison of models for the biogeographical reconstruction computed by the BioGeoBEARS software: dispersal–extinction–cladogenesis model (DEC), dispersal-vicariance analysis (DIVALIKE) and BayArea-like model (BAYAREALIKE) models and their respective alternatives, including the parameter *J*, which represents per-event weight of founder event connected with cladogenesis event. The best model is highlighted **in bold**. Parameters of the models are *d* (dispersal) describing the rate of range expansion, *e* (extinction) mirroring potential range contraction and *j,* which represents founder effect.

| MODEL | LnL | No. params | D | e | J | ΔAICc | AICc weight |
|---|---|---|---|---|---|---|---|
| DEC | -112.3 | 2 | 0.02 | <0.001 | 0 | 12.7 | <0.01 |
| DEC+J | -108.1 | 3 | 0.02 | <0.001 | 0.03 | 6.4 | 0.04 |
| DIVALIKE | -116.5 | 2 | 0.03 | <0.001 | 0 | 21 | <0.01 |
| DIVALIKE+J | -114.5 | 3 | 0.02 | <0.001 | 0.03 | 19.2 | <0.01 |
| BAYAREALIKE | -124.8 | 2 | 0.01 | 0.05 | 0 | 37.7 | <0.01 |
| **BAYAREALIKE+J** | **-104.9** | **3** | **0.01** | **<0.001** | **0.05** | **0** | **0.96** |

**Table S14.** Comparison of mean niche overlap (indicated by Schoener's D values) between allopatric and sympatric sister species pairs of the butterfly genus *Erebia*. Niche overlap was calculated for the total climatic niche based on the PCA of four climate variables, for each variable separately, and for the elevation. Annual precipitation was $\log_{10}$-transformed. A generalised linear model (GLM) with Gaussian error distribution was used to compare the values of niche overlap between sister and non-sister species pairs. The Schoener's D values were available with species with > 5 distribution points, in total the analyses included 16 species.

| | | Allopatric species | | Sympatric species | | | |
|---|---|---|---|---|---|---|---|
| **Niche** | **code** | **mean** | **SE** | **mean** | **SE** | **F** | **P** |
| Total climatic niche | | 0.185 | 0.113 | 0.364 | 0.065 | 1.879 | 0.192 |
| Annual mean temperature | bio1 | 0.306 | 0.123 | 0.531 | 0.071 | 2.514 | 0.135 |
| Temperature seasonality | bio4 | 0.327 | 0.100 | 0.532 | 0.058 | 3.129 | 0.099 |
| Annual precipitation | bio12 | 0.294 | 0.109 | 0.526 | 0.063 | 3.380 | 0.087 |
| Precipitation seasonality | bio15 | 0.398 | 0.158 | 0.451 | 0.091 | 0.086 | 0.774 |
| Elevation | elev | 0.444 | 0.114 | 0.492 | 0.066 | 0.137 | 0.717 |
